# Hyperdiverse archaea near life limits at the polyextreme geothermal Dallol area

**DOI:** 10.1101/658211

**Authors:** Jodie Belilla, David Moreira, Ludwig Jardillier, Guillaume Reboul, Karim Benzerara, José M. López-García, Paola Bertolino, Ana I. López-Archilla, Purificación López-García

## Abstract

Microbial life has adapted to various individual extreme conditions; yet, organisms simultaneously adapted to very low pH, high salt and high temperature are unknown. We combined environmental 16S/18S rRNA-gene metabarcoding, cultural approaches, fluorescence-activated cell sorting, scanning electron microscopy and chemical analyses to study samples along such unique polyextreme gradients in the Dallol-Danakil area (Ethiopia). We identify two physicochemical barriers to life in the presence of surface liquid water defined by: i) high chaotropicity-low water activity in Mg^2+^/Ca^2+^-dominated brines and ii) hyperacidity-salt combinations (pH~0/ NaCl-dominated salt-saturation) When detected, life was dominated by highly diverse ultrasmall archaea widely distributed across phyla with and without previously known halophilic members. We hypothesize that high cytoplasmic K^+^-level was an original archaeal adaptation to hyperthermophily, subsequently exapted during multiple transitions to extreme halophily. We detect active silica encrustment/fossilization of cells but also abiotic biomorphs of varied chemistry. Our work helps circumscribing habitability and calls for cautionary interpretations of morphological biosignatures on Earth and beyond.

---

Microbial life has adapted to so-called extreme values of temperature, pH or salinity, but also to several polyextreme, e.g. hot acidic or salty alkaline, ecosystems^1,2^. Various microbial lineages have been identified in acidic brines in the pH range 1.5-4.5, e.g. in Western Australia^3,4^ and Chile^3^. However, although some acidophilic archaea thrive at pH~0 (*Picrophilus oshimae* grows at an optimal pH of 0.7)^5^ and many halophilic archaea live in hypersaline systems (>30%; NaCl-saturation conditions), organisms adapted simultaneously to very low pH (<1) and high salt, and eventually also high temperature, are not known among cultured prokaryotic species^1^. Are molecular adaptations to these combinations incompatible or (hot) hyperacidic hypersaline environments simply rare and unexplored? The Dallol geothermal dome and its surroundings (Danakil Depression, Afar, Ethiopia) allow to address this question by offering unique polyextreme gradients combining high salt content (33 to >50%; either Mg^2+^/Ca^2+^ or Na^+^(/Fe^2+/3+^)-rich), high temperature (25-110°C) and low pH (≤−1.5 to 6).

Dallol is an up-lifted (~40 m) dome structure located in the North of the Danakil depression (~120 m below-sea-level), a 200 km-long basin within the Afar rift, at the triple junction between the Nubian, Somalian and Arabian Plates^6^. Lying only 30 km north of the hypersaline, hydrothermally-influenced, Lake Assale (Karum) and the Erta Ale volcanic range, Dallol does not display volcanic outcrops but intense degassing and hydrothermalism. These activities are observed on the salt dome and the adjacent Black Mountain and Yellow Lake (Gaet’Ale) areas^6,7^ (Fig. 1a-b). Gas and fluid isotopic measurements indicate that meteoritic waters, notably infiltrating from the high Ethiopian plateau (>2,500 m), interact with an underlying geothermal reservoir (280-370°C)^7,8^. Further interaction of those fluids with the km-thick marine evaporites filling the Danakil depression results in unique combinations of polyextreme conditions and salt chemistries^6,7,9,10^, which have led some authors consider Dallol as a Mars analog^11^.

**Fig. 1.**
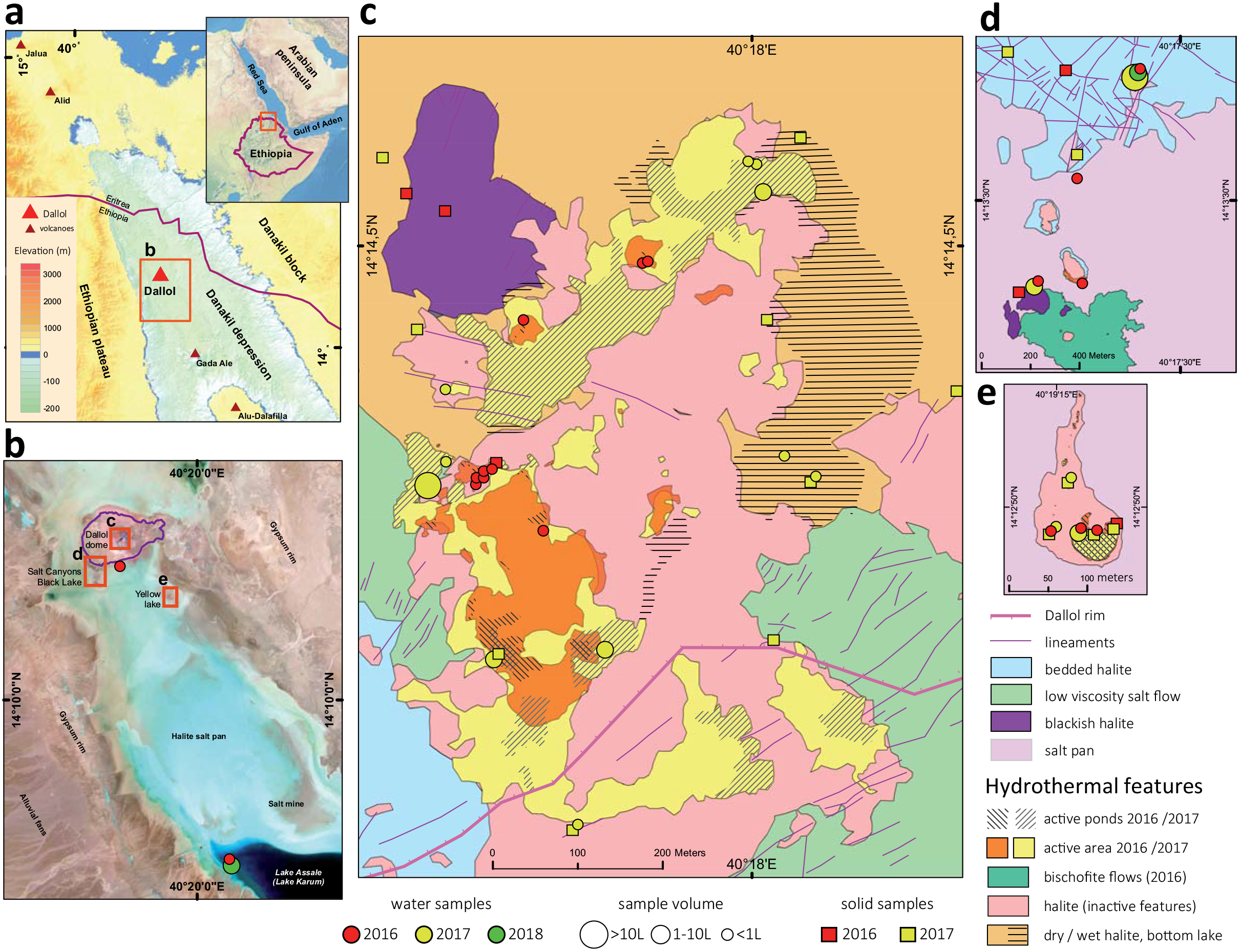
Overview of sampling sites at the polyextreme geothermal field of Dallol and its surroundings in the Danakil Depression, Ethiopia. **a**, Location of the Dallol dome area in the Danakil Depression following the alignment of the Erta Ale volcanic range (Gada Ale, Alu-Dalafilla), Northern Ethiopia; **b**, closer view of the sampling zones in the Dallol area and Lake Assale or Karum (satellite image from Copernicus Sentinel 1; 2017, January 19th); **c-e**, geological maps showing the sampling sites at (**c**) the Dallol dome summit, (**d**) West salt canyons and Black Mountain, including the Black Lake and (**e**) Yellow Lake (Gaet’Ale) zone. Squares and circles indicate the nature of collected samples and their color, the collection date. The size of circles is proportional to the collected brine volume for analyses. Specific sample names are indicated in the aerial view shown in Supplementary Fig. 1.

Here, we use environmental 16S/18S rRNA-gene metabarcoding, cultural approaches, fluorescence-activated cell sorting and scanning electron microscopy combined with chemical analyses to explore microbial occurrence, diversity and potential fossilization along Dallol-Danakil polyextreme gradients^12,13,14,15^.

## Results and Discussion

To investigate the distribution and, eventually, type of microbial life along those polyextreme gradients, we analyzed a large variety of brine and mineral samples collected mainly in two field expeditions (January 2016 and 2017; a few additional samples were collected in 2018) in four major zones (Fig. 1, Supplementary Fig. 1-2, Supplementary Table 1). The first zone corresponded to the hypersaline (37-42%) hyperacidic (pH between ~0 and −1; values down to −1.6 were measured on highly concentrated and oxidized brines on site) and sometimes hot (up to 108°C) colorful ponds on the top of the Dallol dome (Fig. 1c, Supplementary Figs. 1a and 2a-h, Supplementary Table 1). The second zone comprised the salt canyons located at the Southwestern extremity of the Dallol dome and the Black Mountain area that includes the Black Lake (Figs. 1b and 1d; Supplementary Figs. 1b-c and 2l-q). Brine samples collected in a cave reservoir (Gt samples) and in ephemeral pools with varying degrees of geothermal influence at the dome base (PS/PS3) were hypersaline (~35%), with moderate temperature (~30°C) and acidity (pH ~4-6). By contrast, pools located near the small (~15 m diameter), extremely hypersaline (>70%), hot (~70°C) and acidic (pH~3) Black Lake were slightly more acidic (pH~3), warmer (40°C) and hypersaline (35-60%) than dome-base pools (PSBL; Supplementary Table 1). The third zone corresponded to the Yellow Lake and neighboring ponds, an acidic (pH~1.8), warm (~40°C) and extremely hypersaline^16^ system (≥50%) actively emitting toxic gases. These include light hydrocarbons^8^, as attested by numerous dead birds around (Fig. 1e, Supplementary Figs. 1d and 2i-k). The fourth zone comprised the hypersaline (36%), almost neutral (pH~6.5), Lake Assale (Fig. 1b, Supplementary Fig. 2r), which we used as a milder, yet extreme, Danakil system for comparison. In contrast with a continuous degassing activity, the hydrothermal manifestations were highly dynamic, especially on the dome and the Black Mountain area. Indeed, the area affected by hydrothermal activity in January 2017 was much more extensive than the year before (Fig. 1 and Supplementary Fig. 1). Dallol chimneys and hyperacidic ponds can appear and desiccate in a matter of days or weeks, generating a variety of evaporitic crystalline structures observable in situ^17^. Likewise, very active, occasionally explosive (salt ‘bombs’), hydrothermal activity characterized by hot (110°C), slightly acidic (pH~4.4), black hypersaline fluids was detected in the Black Mountain area in 2016 (‘Little Dallol’; sample BL6-01; Supplementary Figs. 1b and 2l) but not in the following years. Also, active bischofite flows^6,7,18^ (116°C) were observed in the Black Mountain area in 2016 but not in 2017.

**Fig. 2.**
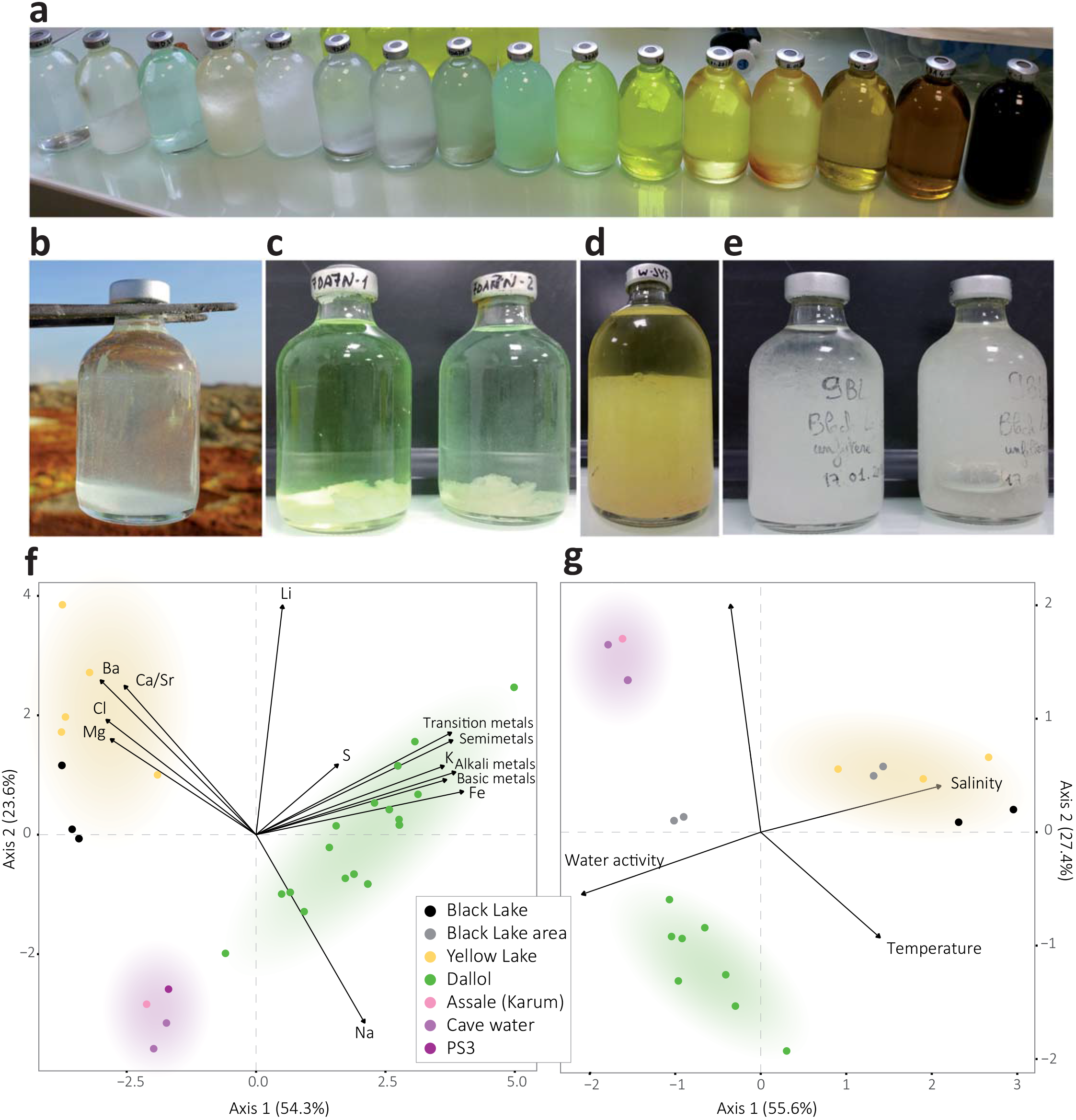
Physicochemical features of liquid samples from the Dallol area. **a**, overview of the color palette showed by samples analyzed in this study, reflecting different chemistries and oxidation states; **b-e**, examples of salt-oversaturated samples; **b**, immediate (seconds) precipitation of halite crystals as water from a hot spring (108°C) cools down upon collection; **c-e**, salt precipitates forming after storage at ca. 8°C in water collected from (**c**) Dallol hyperacidic ponds, (**d**) Yellow Lake and (**e**) Black Lake; **f**, Principal Component Analysis (PCA) of 29 samples according to their chemical composition (see Supplementary Table 2). Transition metals group Cr, Mo, Mn, Sc, Zn, V, U, Ce, La, Cu; semimetals, As, B, Sb, Si; basic metals, Tl, Al, Ga, Sh; and alkali metals, Rb, Cs. Some elements are highlighted out of these groups owing to their high relative abundance or to their distant placement. A PCA showing all individual metal variables can be seen in Supplementary Fig. 3a. **g**, PCA of 21samples and key potentially life-limiting physicochemical parameters in the Dallol area (temperature, pH, salinity (TS), water activity). Water activity and salinity-related parameters are provided in Supplementary Table 3. Colored zones in PCA analyses highlight the clusters of samples; they correspond to the three major chemical zones identified in this study.

To assess potential correlations between microbial life and local chemistry, we analyzed the chemical composition of representative samples used in parallel for microbial diversity analyses (see Methods). Our results revealed three major types of solution chemistry depending on the dominant elements (Fig. 2a; Supplementary Fig. 3a). In agreement with recent observations, Dallol ponds were characterized by NaCl supersaturated brines highly enriched in Fe with different oxidation states, largely explaining color variation^17^. Potassium and sulfur were also abundant (Supplementary Table 2). By contrast, samples from the salt canyons and plain near Dallol and Lake Assale were essentially NaCl-dominated, with much lower Fe content, while the Yellow and Black lakes and associated ponds had very high Mg^2+^ and Ca^2+^ concentrations (Supplementary Table 2). Many aromatic compounds were identified, especially in Dallol and Yellow Lake fluids (Supplementary Data 1). Because high chaotropicity associated with Mg_2_Cl-rich brines, high ionic strength and low water activity (a_w_) are thought to be limiting factors for life^12,13,19,20^, we determined these parameters in representative samples (Supplementary Table 3). Based on our experimental measures and theoretical calculations from dominant salts, only samples in the Black and Yellow lake areas displayed life-limiting chaotropicity and a_w_ values according to established limits^12,13,19,20^. A principal component analysis (PCA) showed that the sampled environments were distributed in three major groups depending on solution chemistry, pH and temperature: Black and Yellow Lake samples, anticorrelating with a_w_; Dallol dome samples, mostly correlating with a_w_ but anticorrelating with pH; and Dallol canyon cave reservoir (Gt samples) and Lake Assale, correlating both with a_w_ and pH (Fig. 2g). These results are consistent with those obtained with ANOVA and subsequent post-hoc analysis, which show significant differences between the majority of the groups among them and for the variables tested (Supplementary Table 4).

**Fig. 3.**
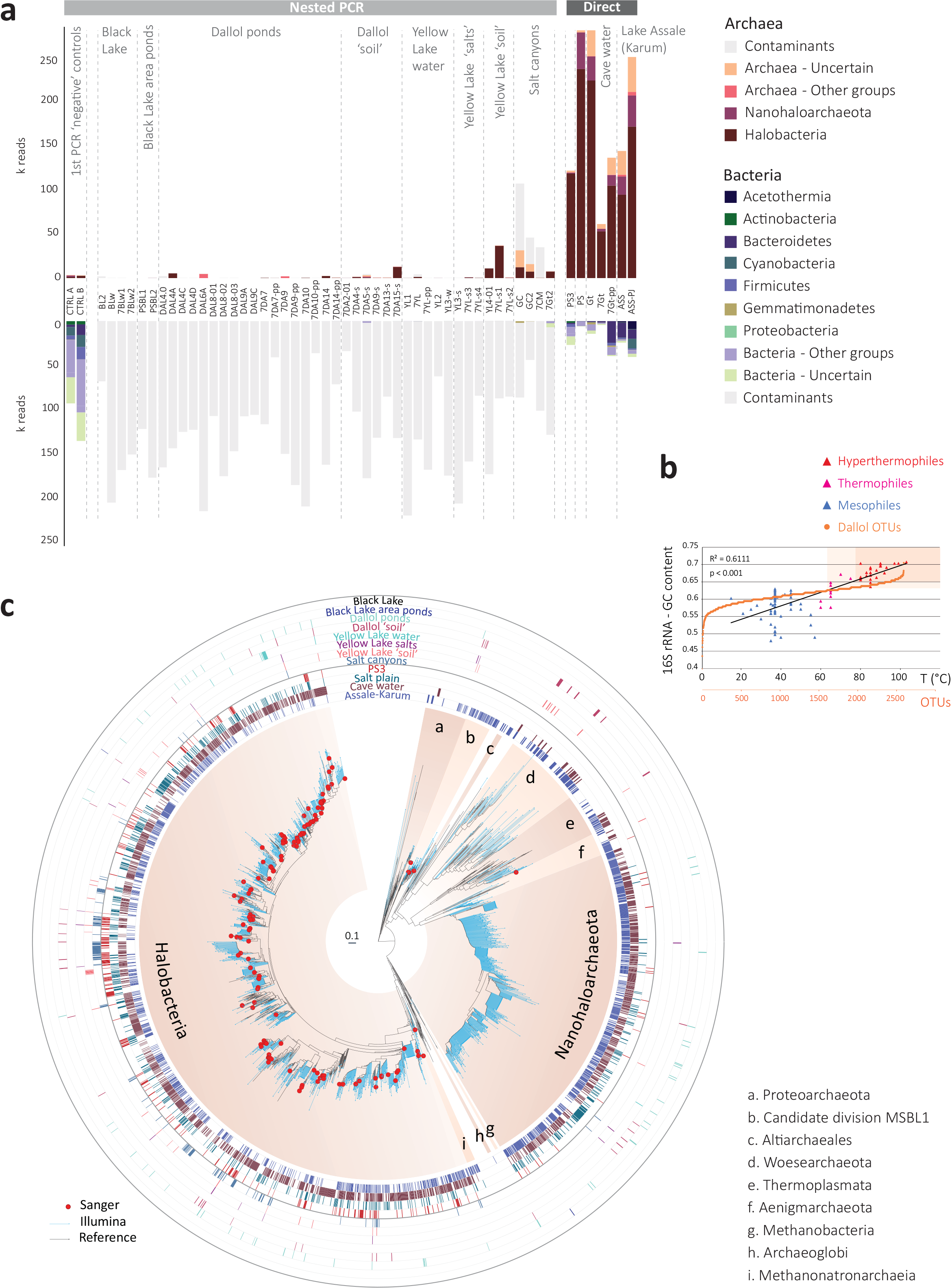
Distribution and diversity of prokaryotes in samples from the Dallol mound and surrounding areas based on 16S rRNA gene metabarcoding data. **a**, histograms showing the presence/absence and abundance of amplicon reads of archaea (upper panel) and bacteria (lower panel) obtained with universal prokaryotic primers. Samples yielding amplicons directly (negative PCR controls were negative) are shown on the right (Direct). Samples for which amplicons were only obtained after nested PCR, all of which also yielded amplicons in ‘negative’ controls, are displayed on the left (Nested PCR). Sequences identified in the ‘negative’ controls, considered as contaminants, are shaded in light grey in the corresponding Dallol samples. The phylogenetic affiliation of dominant archaeal and bacterial groups is color-coded. For details, see Supplementary Data 2-3. **b**, GC content of archaeal OTUs plotted on a graph showing the positive correlation of GC content (for the same 16S rRNA region) and growth temperature of diverse described archaeal species. **c**, phylogenetic tree of archaeal 16S rRNA gene sequences showing the phylogenetic placement of archaeal OTUs identified in the different environmental samples (full tree provided as Supplementary Data 4). Sequences derived from metabarcoding studies are represented with blue branches (Illumina sequences); those derived from cloning and Sanger sequencing of environmental samples, cultures and FACS-sorted cells are labelled with a red dot. Reference sequences are in black. Concentric circles around the tree indicate the presence/absence of the corresponding OTUs in different groups of samples (groups shown in panel a).

To ascertain the occurrence and diversity of microbial life along these physicochemical gradients, we purified DNA from a broad selection of brine samples (0.2-30-μm cell fraction), and solid samples (gypsum and halite-rich salt crusts, compacted sediment and soil-like samples; Supplementary Table 1). We carried out 16S/18S rRNA gene-based diversity studies by high-throughput short-amplicon sequencing (metabarcoding approach) but also sequenced almost-full-length genes from clone libraries, providing local reference sequences for more accurate phylogenetic analyses (see Methods). Despite intensive PCR efforts and extensive sampling in Dallol polyextreme ponds, including pools that were active in two consecutive years (Supplementary Fig. 1) to minimize ephemeral system-derived effects, we only amplified 16S/18S rRNA genes from Dallol canyon cave water, the dome-base geothermally-influenced salt plain and Lake Assale, but never from the Dallol dome and Black/Yellow lakes (Fig. 3a). To check whether this resulted from excessively low DNA amounts in those samples (although relatively large volumes were filtered), we carried out semi-nested PCR reactions using as templates potential amplicons produced during the first PCR-amplification reaction, including first-PCR negative controls. Almost all samples produced amplicons in semi-nested PCR-reactions, including the first-PCR blanks (Fig. 3a). Metabarcoding analysis revealed that amplicons from direct PCR-reactions (PS/PS3, Gt, Assale) were largely dominated by archaeal sequences (>85%), grouping in diverse and abundant OTUs (Supplementary Table 5). By contrast, amplicons derived from Dallol ponds, Black and Yellow lakes but also first-PCR ‘negative’-controls were dominated by bacterial sequences. Most of them were related to well-known kit and laboratory contaminants^21,22^, other were human-related bacteria likely introduced during intensive afar and tourist daily visits to the site; a few archaeal sequences might result from aerosol cross-contamination despite extensive laboratory precautions (see Methods). After removal of contaminant sequences (grey bars, Fig. 3a; Supplementary Data 2), only rare OTUs encompassing few reads (mostly archaeal) could be associated to Dallol dome or Yellow Lake brines, which we interpret as likely dispersal forms (dusty wind is frequent in the area). Slightly higher abundances of archaeal OTUs were identified in ‘soil’ samples, i.e. samples retrieved from salty consolidated mud or crusts where dust brought by the wind from the surrounding plateaus accumulates and starts constituting a proto-soil (with incipient microbial communities; e.g. Supplementary Fig. 2i). Therefore, while we cannot exclude the presence of active life in these ‘soil’ samples, our results strongly suggest that active microbial life is absent from polyextreme Dallol ponds and the Black and Yellow lakes.

By contrast, PS/PS3, Gt and Assale samples harbored extremely diverse archaea (2,653 OTUs conservatively determined at 95% identity, i.e. genus level) that virtually spanned the known archaeal diversity (Fig. 3; Supplementary Table 5; Supplementary Data 2). Around half of that diversity belonged to Halobacteria, and an additional quarter to the Nanohaloarchaeota^23^. The rest of archaea distributed in lineages typically present in hypersaline environments, e.g. the Methanonatronoarchaeia^24,25^ and Candidate Divison MSBL1, thought to encompass methanogens^26^ and/or sugar-fermentors^27^, but also other archaeal groups not specifically associated with salty systems (although can sometimes be detected in hypersaline settings, e.g. some Thermoplasmata or Woesearchaeota). These included Thermoplasmata and Archaeoglobi within Euryarchaeota, Woesearchaeota and other lineages (Aenigmarchaeota, Altiarchaeales) usually grouped as DPANN^28–30^ and Thaumarchaeota and Crenarchaeota (Sulfolobales) within the TACK/Proteoarchaeota^31^ (Fig. 3a; Supplementary Data 2). In addition, based on the fact that rRNA GC content correlates with growth temperature, around 27% and 6% of archaeal OTUs were inferred to correspond to, respectively, thermophilic and hyperthermophilic organisms (see Methods; Fig. 3b). As previously observed^23,28,29^, common archaeal primers for near-full 16S rRNA genes (Fig. 3c, red dots) failed to amplify Nanohaloarchaeota and other divergent DPANN lineages. These likely encompass ectosymbionts/parasites^28–30,32^. Given their relative abundance and co-occurrence in these and other ecosystems, it is tempting to hypothesize that Nanohaloarchaeota are (ecto)symbionts of Halobacteria; likewise, Woesearchaeota might potentially be associated with *Thermoplasma*-like archaea. Although much less abundant, bacteria belonging to diverse phyla, including CPR (Candidate Phyla Radiation) lineages, were also present in these samples (710 OTUs; Supplementary Fig. 4; Supplementary Table 5; Supplementary Data 2). In addition to typical extreme halophilic genera (e.g. *Salinibacter*, Bacteroidetes), one Deltaproteobacteria group and two divergent bacterial clades were overrepresented in Dallol canyon Gt samples. Less abundant and diverse, eukaryotes were present in Lake Assale and, occasionally, the salt plain and Gt, being dominated by halophilic *Dunaliella* algae (Supplementary Fig. 5; Supplementary Data 3).

**Fig. 4.**
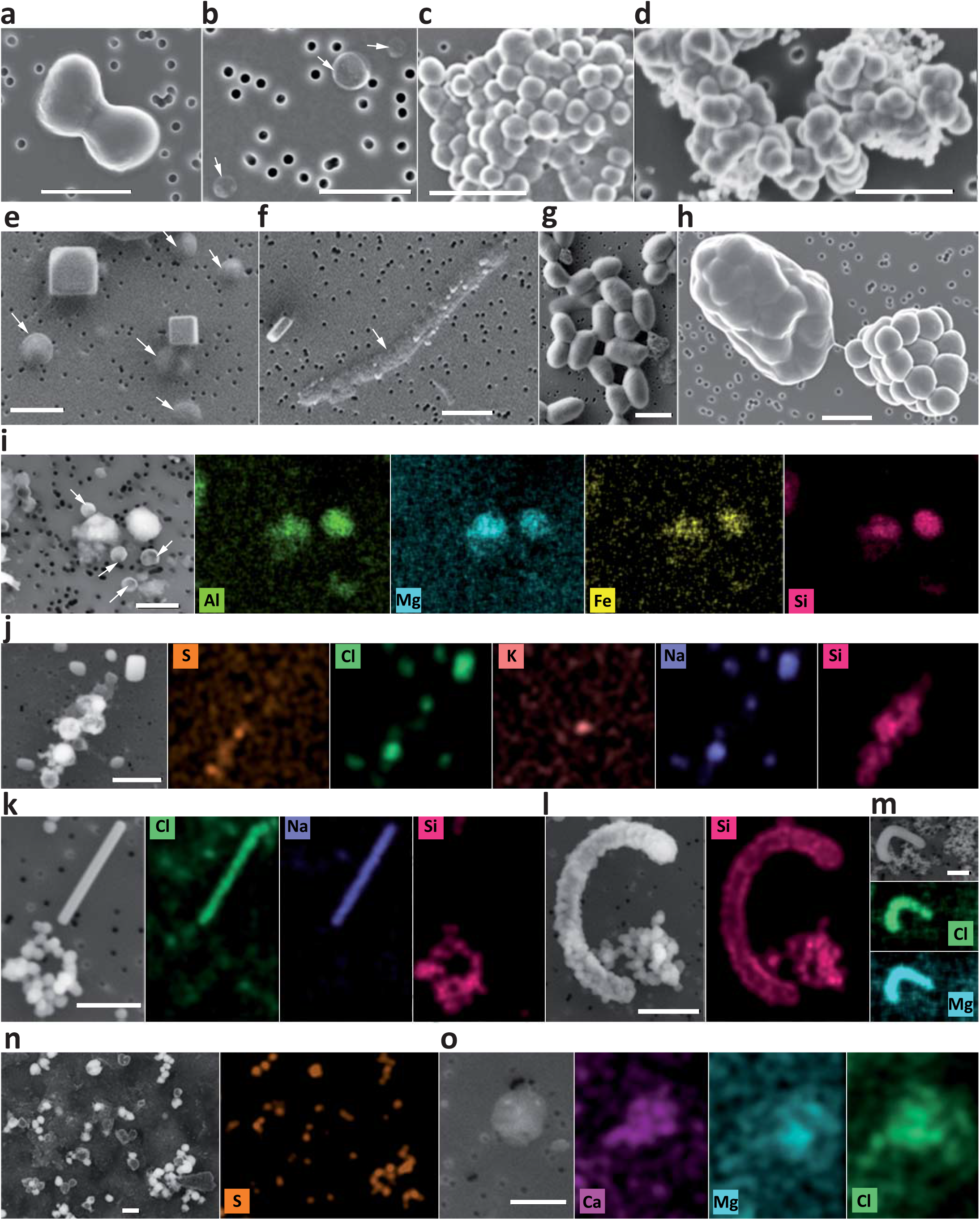
Scanning electron microscopy (SEM) pictures and chemical maps of cells and abiotic biomorphs identified in samples from the Dallol region. **a-h**, SEM pictures of cells (**a-c**, **e-h**) and abiotic biomorphs (**d**). i-o, SEM images and associated chemical maps of cells and biomorphs; color intensity provides semi-quantitative information of the mapped elements. **a**, FACS-sorted dividing cells from sample PS (hydrated salt pan between the Dallol dome base and the Black Lake); **b**, FACS-sorted ultrasmall cells from 7Gt samples (cave water reservoir, Dallol canyons); **c**, FACS-sorted colony of ultrasmall cells from sample PS (note cytoplasmic bridges between cells); **d**, FACS-sorted abiotic silica biomorphs from the Dallol pond 7DA9 (note the similar shape and morphology as compared to cells in panel c); **e**, cocci and halite crystals in 8Gt samples (cave water); **f**, long rod in 8Gt; **g**, FACS-sorted cells from Gt samples; **h**, FACS-sorted colonies from sample PS (note the bridge between one naked colony and one colony covered by an exopolymeric-like matrix); **i**, small cocci and amorphous Al-Mg-Fe-rich silicate minerals from Gt; **j**, NaCl crystals and S-Si-rich abiotic biomorphs from Dallol pond sample 7DA7; **k**, NaCl crystal and Si-biomorphs and **l**, Si-encrusted cell and Si-biomorphs in sample 8Ass (Lake Assale); **m**, Mg-Cl biomorph in sample BLPS_04 (Black Lake area pond); **n**, S-rich biomorphs in Dallol pond 7DA9; **o**, Ca-Mg-Cl biomorph in YL-w2 (Yellow Lake pond). SEM photographs were taken using In Lens or AsB detectors; AsB was used for chemical mapping purposes. For additional images and SEM details, see Supplementary Data 6-7. White arrows indicate cells difficult to recognize due to their small size and/or flattened aspect possibly resulting from sample preparation and/or high vacuum conditions within the SEM. The scale bar corresponds to 1 μm.

Consistent with metabarcoding results, and despite the use of various culture media and growth conditions mimicking local environments (see Methods), cultural approaches did not yield enrichments for any of the Dallol dome, Black and Yellow lake samples. We easily obtained enrichments from the canyon cave (Gt/7Gt) and salt plain (PS/PS3) samples in most culture media (except on benzoate/hexadecane) and tested conditions (except at 70°C in the dark). However, all attempts to isolate microorganisms at pH<3 from these enrichments also failed. The most acidophilic isolate obtained from serial dilutions (PS3-A1) grew only at 37°C and optimal pH 5.5 (range 3-7). Its 16S rRNA gene was 98.5% identical to that of *Halarchaeum rubridurum* MH1-16-3 (NR_112764), an acidophilic haloarchaeon growing at pH 4.0-6.5^33^. In agreement with metabarcoding and culture-derived observations, multiparametric fluorescence analysis showed no DNA fluorescence above background for Dallol and Yellow Lake samples (Supplementary Fig. 6). Because optical and scanning electron microscopy (SEM) observations suggested that indigenous cells were unusually small, we applied fluorescence-activated cell-sorting (FACS) to samples from the different Dallol environments, including samples with almost no events above noise (Supplementary Table 1) followed by systematic SEM analysis of sorted events. We only detected cells in Dallol cave water and salt plain samples but not in dome ponds or Yellow Lake samples (Supplementary Fig. 6). Consistent with this, after DNA purification of FACS-sorted particles, 16S rRNA amplicons could only be obtained from different cave and salt plain samples but not from Dallol dome or Yellow Lake samples. Cell counts estimated from FACS for the cave and salt plain samples were low (average 3.1 × 10^4^ cells.ml^−1^ and 5.3 × 10^4^ cells.ml^−1^ for the cave and PS samples, respectively). Sorted cells were usually small to ultrasmall (down to 0.25-0.3 μm diameter; Fig. 4). In PS samples, some of these small cells formed colonies (Supplementary Fig. 6, Fig. 4c) sometimes surrounded by an exopolymeric matrix cover (Fig. 4h). The presence of cytoplasmic bridges and/or potential cell fusions (Supplementary Fig. 6, Fig. 4c) suggest that they might be archaeal colonies^34^. FACS-sorted fluorescent particles in Dallol pond samples appeared to correspond exclusively to salt crystals or cell-sized amorphous minerals morphologically resembling cells, i.e. biomorphs^35,36^ (e.g. Fig. 4d in comparison with Fig. 4c). This prompted us to carry out a more systematic search for abiotic biomorphs in our samples. SEM observations coupled with chemical mapping by energy dispersive X-ray spectrometry (EDXS) showed a variety of cocci-like biomorph structures of diverse elemental compositions. Many of them were Si biomorphs (Dallol ponds, Yellow and Assale lakes), but we also detected Fe-Al silicates (Gt), S or S-rich biomorphs (Dallol ponds), and Ca or Mg chlorides (Yellow lake, BLPS samples). (Fig. 4; Supplementary Table 6; Supplementary Data 6-7). At the same time, we observed Si-encrusted rod-shaped cells in Lake Assale samples (Fig. 4.l). Therefore, silica rounded precipitates represent ultrasmall cell-like biomorphs in samples with no detectable life but contribute to cell encrustment and potential fossilization when life is present.

Our work has three major implications. First, by studying the microbial distribution along gradients of polyextreme conditions in the geothermal area of Dallol and its surroundings in the Danakil Depression, we identify two major physicochemical barriers that prevent life to thrive in the presence of liquid water on the surface of our planet and, potentially, elsewhere^14^, despite it is a widely accepted criterion for habitability. Confirming previous studies^12,13,19,20^, one such barrier is imposed by high chaotropicity and low a_w_, which are associated to high Mg^2+^-brines in Black and Yellow lake areas. The second barrier seems to be imposed by the hyperacid-hypersaline combinations found in the Dallol dome ponds (pH~0; salt>35%), regardless of temperature. This suggests that molecular adaptations to simultaneous very low-pH and high-salt extremes are incompatible beyond those limits. In principle, more acidic proteins, intracellular K^+^ accumulation (‘salt-in’ strategy) or internal positive membrane potential generated by cations or H^+^/cation antiporters serve both acidophilic and halophilic adaptations^37–39^. However, membrane stability/function problems and/or high external Cl^−^ concentrations inducing H^+^ and cation (K^+^/Na^+^) import and potentially disrupting membrane bioenergetics^38^, might be deleterious under these conditions. We cannot exclude other explanations linked to the presence of multiple stressors, such as high metal content, or an increased susceptibility to the presence of local chaotropic salts in the Dallol hyperacidic ponds even if measured chaotropicity values are relatively low (−31 to +19 kJ/kg) as compared to the established limit for life (87.3 kJ/kg)^12,13,20^ (Supplementary Table 3). Future studies should help to identify the molecular barriers limiting the adaptation of life to this combination of extremes. Second, although extreme environments usually are low-diversity systems, we identify here exceptionally diverse and abundant archaea spanning known major taxa in hypersaline, mildly acidic systems near life-limit conditions. A wide archaeal (and to a lesser extent, bacterial) diversity seems consistent with suggestions that NaCl-dominated brines are not as extreme as previously thought^40^ but also with recent observations that the mixing of meteoric and geothermal fluids leads to hyperdiverse communities^41^. Nonetheless, life at high salt requires extensive molecular adaptations^12,13,19,40^, which might seem at odds with multiple independent adaptations to extreme halophily across archaea. Among those adaptations, the intracellular accumulation of K^+^ (‘salt-in’ strategy), accompanied by the corresponding adaptation of intracellular proteins to function under those conditions, has been crucial. Based on the observation that the deepest archaeal branches correspond to (hyper)thermophilic lineages^42^ and that non-halophilic hyperthermophilic archaea accumulate high intracellular K^+^ (1.1-3M) for protein thermoprotection^43,44^ (thermoacidophiles also need K^+^ for pH homeostasis^38^), we hypothesize that intracellular K^+^ accumulation is an ancestral archaeal trait that has been independently exapted in different taxa for adaptation to hypersaline habitats. Finally, the extensive occurrence of abiotic, mostly Si-rich, biomorphs mimicking the simple shape and size of ultrasmall cells in the hydrothermally-influenced Dallol settings reinforces the equivocal nature of morphological microfossils^35^ and calls for the combination of multiple biosignatures before claiming the presence of life on the early Earth and beyond.

## Materials and Methods

### Sampling and measurement of physicochemical parameters on site

Samples were collected during two field trips carried out in January 2016 and January 2017 (when air temperature rarely exceeded 40-45°C); a few additional samples were collected in January 2018 (Fig. 1; Supplementary Fig. 1 and Supplementary Table 1). All sampling points and mapping data were georeferenced using a Trimble^®^ handheld GPS (Juno SB series) equipped with ESRI software ArcPad^®^ 10. Cartography of hydrogeothermal activity areas was generated using ESRI GIS ArcMap™ mapping software ArcGis^®^ 10.1 over georeferenced Phantom-4 drone images taken by O. Grunewald during field campaigns, compared with and updating previous local geological cartography^7^. Samples were collected in three major areas at the Dallol dome and its vicinity (Fig. 1b): i) the top of the Dallol dome, comprising various hydrothermal pools with diverse degrees of oxidation (Fig.1c); ii) the Black Mountain area (Fig. 1d), including the Black Lake and surrounding bischofite flows and the South-Western salt canyons harboring water reservoirs often influenced by the geothermal activity and iii) the Yellow Lake (Gaet’Ale) area (Fig. 1e). We also collected samples from the hypersaline Lake Assale (Karum), located a few kilometers to the South in the Danakil Depression (Fig. 1b). Physicochemical parameters (Supplementary Table 1) were measured in situ with a YSI Professional Series Plus multiparameter probe (pH, temperature, dissolved oxygen, redox potential) up to 70°C and a Hanna HI93530 temperature probe (working range −200/1,000°C) and a Hanna HI991001 pH probe (working pH range −2.00/16.00) at higher temperatures. Salinity was measured in situ with a refractometer on 1:10 dilutions in MilliQ water. Brine samples for chemical analyses were collected in 50 ml glass bottles after prefiltration through 0.22 μm pore-diameter filters; bottles were filled to the top and sealed with rubber stoppers to prevent the (further) oxidation of reduced fluids. Solid and water samples for microbial diversity analyses and culturing assays were collected under the most possible aseptic conditions to prevent contamination (gloves, sterile forceps and containers). Samples for culture assays were kept at room temperature. Salts and mineral fragments for DNA-based analyses were conditioned in Falcon tubes and fixed with absolute ethanol. Water samples (volumes for each sample are indicated in Supplementary Table 1) were filtered through 30 μm pore-diameter filters to remove large particles and sequentially filtered either through 0.22 μm pore-diameter filters (Whatman^®^) or using 0.2 μm pore-size Cell-Trap units (MEM-TEQ Ventures Ltd, Wigan, UK). Filters or Cell-Trap concentrates retaining 0.2-30 μm particles were fixed in 2-ml cryotubes with absolute ethanol (>80% final concentration). Back in the laboratory, ethanol-fixed samples were stored at −20°C until use.

### Chemical analyses, salinity, chaotropicity, ionic strength and water activity

The chemical composition of solid and 0.2 μm-prefiltered liquid samples was analyzed at the SIDI service (Universidad Autónoma de Madrid). Major and trace elements in liquid samples were analyzed by total reflection X-ray fluorescence (TXRF) with a TXRF-8030c FEI spectrometer and inductively coupled plasma mass spectrometry (ICP-MS) using a Perkin-Elmer NexION 300XX instrument. Ions were analyzed using a Dionex DX-600 ion chromatography system. Organic molecules were characterized using a Varian HPLC-DAD/FL/LS liquid chromatograph. Crystalline phases in solid samples were characterized by x-ray diffraction using a X’Pert PRO Theta/Theta diffractometer (Panalytical) and identified by comparison with the International Centre for Diffraction Data (ICDD) PDF-4+ database using the ‘High Score Plus’ software (Malvern Panalytical https://www.malvernpanalytical.com/es/products/category/software/x-ray-diffraction-software/highscore-with-plus-option). Inorganic data are provided in Supplementary Table 2; organic and ionic chemistry data in Supplementary Data 1. Salinity (weight/volume) was also experimentally measured in triplicates (and up to 6 replicates) by weighting the total solids after heat-drying 1 ml aliquots in ceramic crucibles at 120°C for at least 24h. Chaotropicity was experimentally measured according to the temperature of gelation of ultrapure gelatin (for Ca-rich samples) and agar (rest of samples) determined using the spectrometric assay developed by Cray et al.^45^ (Supplementary Table 3). Chaotropicity was also calculated according to Cray and coworkers^46^ based on the abundance of dominant Na, K, Mg, Ca and Fe cations and, on the ground that Cl is the dominant anion, assuming they essentially form chlorine salts (NaCl, KCl, MgCl_2_, CaCl_2_ and FeCl_2_). Ionic strength was calculated according to Fox-Powell et al.^47^. Water activity was measured on 10-ml unfiltered aliquots at room temperature using a HC2-AW probe and HP23-AW-A indicator (Rotronic AG, Bassersdorf, Switzerland) calibrated at 23°C using the AwQuick acquisition mode (error per measure 0.0027). Principal component analyses (PCA) of samples, chemical and physicochemical parameters (Fig. 2 and Supplementary Fig. 3) were done using R-software^48^ packages FactoMineR^49^ and factoextra^50^. Differences between the groups of samples belonging to the same physicochemical zone segregating in the PCA were tested using the one-way ANOVA module of IBM SPSS Statistics 24 software. The significance of differences among groups and with the measured parameters were checked by means of a post-hoc comparison using the Bonferroni test.

### DNA purification and 16/18S rRNA gene metabarcoding

DNA from filters, Cell-Trap concentrates and grinded solid samples was purified using the Power Soil DNA Isolation Kit (MoBio, Carlsbad, CA, USA) under a UV-irradiated Erlab CaptairBio DNA/RNA PCR Workstation. Prior to DNA purification, filters were cut in small pieces with a sterile scalpel and ethanol remaining in cryotubes filtered through 0.2 μm pore-diameter filters and processed in the same way. Ethanol-fixed Cell-Trap concentrates were centrifuged for 10 min at 13,000 rpm and the pellet resuspended in the first kit buffer. Samples were let rehydrate for at least 2h at 4°C in the kit resuspension buffer. For a selection of Cell-Trap concentrates, FACS-sorted cells and to monitor potential culture enrichments, we also used the Arcturus PicoPure DNA Isolation kit (Applied Biosystems – Foster City, CA, USA; samples labeled pp). DNA was resuspended in 10 mM Tris-HCl, pH 8.0 and stored at −20°C. Bacterial and archaeal 16S rRNA gene fragments of approximatively 290 bp encompassing the V4 hypervariable region were PCR-amplified using U515F (5’-GTGCCAGCMGCCGCGGTAA) and U806R (5’-GGACTACVSGGGTATCTAAT) primers. PCR reactions were conducted in 25 μl, using 1.5 mM MgCl_2,_ 0.2 mM of each dNTP (PCR Nucleotide Mix, Promega), 0.1 μM of each primer, 1 to 5 μl of purified ‘DNA’ and 1 U of the hot-start Taq Platinum polymerase (Invitrogen, Carlsbad, CA, USA). GoTaq (Promega) was also tried when amplicons were not detected but did not yield better results. Amplification reactions proceeded for 35 cycles (94°C for 15 s, 50 to 55°C for 30 s and 72°C for 90 s), after a 2 min-denaturation step at 94°C and before a final extension at 72°C for 10 min. Amplicons were visualized after gel electrophoresis and ultrasensitive GelRed^®^ nucleic acid gel stain (Biotium, Fremont, CA, USA) on a UV-light transilluminator. When direct PCR reactions failed to yield amplicons after several assays, PCR conditions and using increasing amounts of input potential DNA, semi-nested reactions using those primers were carried out using as template 1 μl of PCR products, including negative controls, from a first amplification reaction done with universal prokaryotic primers U340F (5’-CCTACGGGRBGCASCAG) and U806R. Eukaryotic 18S rRNA gene fragments including the V4 hypervariable region were amplified using primers EK-565F (5’-GCAGTTAAAAAGCTCGTAGT) and 18S-EUK-1134-R-UNonMet (5’-TTTAAGTTTCAGCCTTGCG). Primers were tagged with different Molecular IDentifiers (MIDs) to allow multiplexing and subsequent sequence sorting. Amplicons from at least 5 independent PCR products for each sample were pooled together and then purified using the QIAquick PCR purification kit (Qiagen, Hilden, Germany). Whenever semi-nested PCR reactions yielded amplicons, semi-nested reactions using as input first-PCR negative controls also yielded amplicons (second-PCR controls did not yield amplicons). Products of these positive ‘negative’ controls were pooled in two control sets (1 and 2) and sequenced along with the rest of amplicons. DNA concentrations were measured using Qubit™ dsDNA HS assays (Invitrogen). Equivalent amplicon amounts obtained for 54 samples (including controls) were multiplexed and sequenced using paired-end (2×300 bp) MiSeq Illumina technology (Eurofins Genomics, Ebersberg, Germany). In parallel, we tried to amplify near-complete 16S/18S rRNA gene fragments (~1400-1500 bp) using combinations of forward archaea-specific primers (21F, 5’-TTCCGGTTGATCCTGCCGGA; Ar109F, 5’-ACKGCTGCTCAGTAACACGT) and bacteria-specific primers (27F, 5’-AGAGTTTGATCCTGGCTCAG) with the prokaryotic reverse primer 1492R (5’-GGTTACCTTGTTACGACTT) and eukaryotic primers 82F (5’-GAAACTGCGAATGGCTC) and 1520R (5’-CYGCAGGTTCACCTAC). When amplified, DNA fragments were cloned using TopoTA™ cloning (Invitrogen) and clone inserts were Sanger-sequenced to yield longer reference sequences. Forward and reverse Sanger sequences were quality-controlled and merged using Codon Code Aligner (http://www.codoncode.com/aligner/).

### Sequence treatment and phylogenetic analyses

Paired-end reads were merged and treated using a combination of existing software to check quality, eliminate primers and MIDs and eliminate potential chimeras. Sequence statistics are given in Supplementary Table 5. Briefly, read merging was done with FLASH^51^, primers and MIDs trimmed with cutadapt^52^ and clean merged reads dereplicated using vsearch^53^, with the uchime_denovo option to eliminate potential chimeras. The resulting dereplicated clean merged reads were used then to define operational taxonomic units (OTUs) at 95% identity cut-off using CD-HIT-EST^54^. This cut-off offered i) a reasonable operational approximation to the genus level diversity while producing a manageable number of OTUs to be included in phylogenetic trees (see below) and ii) allowed us a conservative identification of potential contaminants in our semi-nested PCR-derived datasets. Diversity (Simpson), richness (Chao1) and evenness indices were determined using R-package “vegan” (Supplementary Table 5). OTUs were assigned to known taxonomic groups based on similarity with sequences of a local database including sequences from cultured organisms and environmental surveys retrieved from SILVAv128^55^ and PR2v4^56^. The taxonomic assignation of bacteria and archaea was refined by phylogenetic placement of OTU representative sequences in reference phylogenetic trees. To build these trees, we produced, using Mafft-linsi v7.38^57^, alignments of near full-length archaeal and bacterial 16S rRNA gene sequences comprising Sanger sequences from our gene libraries (144 archaeal, 91 bacterial) and selected references for major identified taxa plus the closest blast-hits to our OTUs (702 archaea, 2,922 bacterial). Poorly aligned regions were removed using TrimAl^58^. Maximum likelihood phylogenetic trees were constructed with IQ-TREE^59^ using the GTR model of sequence evolution with a gamma law and taking into account invariable sites (GTR+G+I). Node support was estimated by ultrafast bootstrapping as implemented in IQ-TREE. Shorter OTU representative sequences (2,653 archaeal, 710 bacterial) were then added to the reference alignment using MAFFT (accurate -linsi ‘addfragments’ option). This final alignment was split in two files (references and OTUs) before using the EPA-ng tool (https://github.com/Pbdas/epa-ng) to place OTUs in the reference trees reconstructed with IQ-TREE. The jplace files generated by EPA-ng were transformed into newick tree files with the genesis library (https://github.com/lczech/genesis). Tree visualization and ring addition were done with GraphLan^60^. To see whether our OTUs might correspond to thermophilic species, we first plotted the GC content of the 16S rRNA gene region used for metabarcoding analyses of a selection of 88 described archaeal species with optimal growth temperatures ranging from 15 to 103°C. These included representatives of all Halobacteria genera, since they are often characterized by high GC content. A regression analysis confirmed the occurrence of a positive correlation^61^ between rRNA GC content and optimal growth temperature also for this shorter 16S rRNA gene amplified region (Fig. 3b). We then plotted the GC content of our archaeal OTUs on the same graph. Dots corresponding to Halobacteria genera remain out of the dark shadowed area in Fig. 3b.

### Cultures

Parallel culture attempts were carried out in two different laboratories (Orsay and Madrid). We used several culture media derived from a classical halophile’s base mineral growth medium^62^ containing (gl^−1^): NaCl (234), KCl (6), NH_4_Cl (0.5), K_2_HPO_4_ (0.5), (NH_4_)_2_SO_4_ (1), MgSO_4_.7H_2_O (30.5), MnCl_2_.7H_2_O (19.5), CaCl_2_.6H_2_O (1.1) and Na_2_CO_3_ (0.2). pH was adjusted to 4 and 2 with 10N H_2_SO_4_. The autoclaved medium was amended with filter-sterilized cyanocobalamin (1 μM final concentration) and 5 ml of an autoclaved CaCl_2_.6H_2_O 1M stock solution. Medium MDH2 contained yeast extract (1 gl^−1^) and glucose (0.5 gl^−1^). Medium MDSH1 had only 2/3 of each base medium salt concentration plus FeCl_3_ (0.1 gl^−1^) and 10 ml.l^−1^ of Allen’s trace solution. It was supplemented with three energy sources (prepared in 10 ml distilled water at pH2 and sterilized by filtration): yeast extract (1 gl^−1^) and glucose (0.5 gl^−1^) (MDS1-org medium); Na_2_S_2_O_3_ (5 gl^−1^) (MDS1-thio medium) and FeSO_4_.7H_2_O (30 gl^−1^) (MDS1-Fe medium). Medium MDSH2 mimicked more closely some Dallol salts as it also contained (gl^−1^): FeCl_3_ (0.1), MnCl_2_.4H_2_O (0.7), CuSO_4_ (0.02), ZnSO_4_.7H_2_O (0.05) and LiCl (0.2) as well as 10 ml l^−1^ of Allen’s trace solution combined with the same energy sources used for MDSH1, yielding media MDSH2-org, MDSH2-thio and MDSH2-Fe. For enrichment cultures, we added 0.1 ml liquid samples to 5 ml medium at pH 2 and 4 and incubated at 37, 50 and 70°C in 10-ml sterile glass tubes depending on the original sample temperatures. Three additional variants of the base salt medium supplemented with FeCl_3_ and trace minerals contained 0.2 gl^−1^ yeast extract (SALT-YE), 0.5 gl^−1^ thiosulfate (SALT-THIO) or 0.6 gl^−1^ benzoate and 5 mM hexadecane (SALT-BH). The pH of these media was adjusted with 34% HCl to 1.5 for Dallol and Black Lake samples, and to 3.5 for YL, PS3 and PSBL samples. 1 ml of sample was added to 4 ml of medium and incubated at 45°C in a light regime and at 37 and 70°C in the dark. We also tried cultures in anaerobic conditions. Potential growth was monitored by optical microscopy and, for some samples, SEM. In the rare cases where enrichments were obtained, we attempted isolation by serial dilutions.

### Flow cytometry and fluorescence-activated cell sorting (FACS)

The presence of cell/particle populations above background level in Dallol samples was assessed with a flow-cytometer cell-sorter FACSAria™III (Becton Dickinson). Several DNA dyes were tested for lowest background signal in forward scatter (FSC) red (695±20 nm) and green (530±15 nm) fluorescence (Supplementary Fig. 6a) using sterile SALT-YE medium as blank. DRAQ5™ and SYTO13^®^ (ThermoFisher) were retained and used at 5 μM final concentration to stain samples in the dark at room temperature for 1 h. Cell-Trap concentrated samples were diluted at 20% with 0.1-μm filtered and autoclaved MilliQ^®^ water. The FACSAria™III was set at purity sort mode triggering on the forward scatter (FSC). Fluorescent target cells/particles were gated based on the FSC and red or green fluorescence (Supplementary Fig. 6b) and flow-sorted at a rate of 1-1,000 particles per second. Sorting was conducted using the FACSDiva™ software (Becton Dickinson); figures were done with the FCSExpress 6 software (De Novo Software). Sorted cells/particles were subsequently observed by scanning electron microscopy for characterization. Minimum and maximum cell abundances were estimated based on the number of sorted particles, duration of sorting and minimal (10μl min-1) and maximal (80μl min-1) flow rates of the FACSAria (Becton Dickinson FACSAria manual).

### Scanning electron microscopy (SEM) and elemental analysis

SEM analyses were carried out on natural samples, FACS-sorted cells/particles and a selection of culture attempts. Liquid samples were deposited on top of 0.1 μm pore-diameter filters (Whatman^®^) under a mild vacuum aspiration regime and briefly rinsed with 0.1-μm filtered and autoclaved MilliQ^®^ water under the same vacuum regime. Filters were let dry and sputtered with carbon prior to SEM observations. SEM analyses were performed using a Zeiss ultra55 field emission gun (FEG) SEM. Secondary electron (SE2) images were acquired using an In Lens detector at an accelerating voltage of 2.0 kV and a working distance of 7.5 mm. Backscattered electron images were acquired for chemical mapping using an angle selective backscattered (AsB) detector at an accelerating voltage of 15 kV and a working distance of ~ 7.5 mm. Elemental maps were generated from hyperspectral images (HyperMap) by energy dispersive X-ray spectrometry (EDXS) using an EDS QUANTAX detector. EDXS data were analyzed using the ESPRIT software package (Bruker).

## Data availability

Sanger sequences have been deposited in GenBank (NCBI) with accession numbers MK894601-MK894820 and Illumina sequences in GenBank Short Read Archive with BioProject number PRJNA541281.

## Acknowledgments

We are grateful to Olivier Grunewald for co-organizing the Dallol expeditions, documenting field research and providing drone images and to Jean-Marie Hullot (in memoriam), Françoise Brenckmann and the Fondation Iris for funding the first field trip. We thank Luigi Cantamessa for the in situ logistics and discussions about local history. We acknowledge Dr. Makonen Tafari (Mekelle University), Abdul Ahmed Aliyu and the Afar authorities for local assistance as well as the Ethiopian army and the Afar police for providing security. We thank Jacques Barthélémy, Elektra Kotopoulou and Juanma Garcia-Ruiz for help and discussions during field trips. We thank Hélène Timpano and the UNICELL platform for cell sorting, Ana Gutiérrez-Preciado for bioinformatic assistance, Adrienne Kish and Charly Faveau for allowing us to measure water activity of selected samples at the Muséum National d’Histoire Naturelle, Eric Viollier for discussion on chemical analyses, Corentin Gille for help with cultures and Georis Billo for script help to treat SEM pictures. This research was funded by the CNRS basic annual funding, the CNRS program TELLUS INTERRVIE and the European Research Council under the European Union’s Seventh Framework Program (ERC Grant Agreement 322669). We thank the European COST Action TD1308 ‘Origins’ for funding a short stay of A.L.A. in Orsay. J.B. was financed by the French Ministry of National Education, Research and Technology.

## Author contributions

P.L.G. and D.M. designed and supervised the research. P.L.G. organized the scientific expeditions. J.B., P.L.G., D.M., L.J. and J.M.L.G. collected samples and took measurements in situ. J.B., PL.G. and P.B. carried out molecular biology analyses. J.B., A.L.A. and D.M. performed culture, chemistry analyses and water-salt-related measurements. A.L.A. and J.B. performed statistical analyses. J.B., G.R. and D.M. analyzed metabarcoding data. K.B. performed SEM and EDX analyses. J.M.L.G. mapped geothermal activity and georeferenced all samples. L.J. and J.B. performed FACS-derived analyses. P.L.G. and J.B. wrote the manuscript. All authors read and commented on the manuscript.

## Supplementary Materials

Supplementary Figures 1-6

Supplementary Tables 1-6

Supplementary Data 1 to 7

## References

1 Harrison, J. P., Gheeraert, N., Tsigelnitskiy, D. & Cockell, C. S. The limits for life under multiple extremes. Trends in microbiology 21, 204–212 (2013).

2 Merino, N. et al. Living at the extremes: Extremophiles and the limits of life in a planetary context. Frontiers in microbiology 10(2019).

3 Johnson, S. S., Chevrette, M. G., Ehlmann, B. L. & Benison, K. C. Insights from the metagenome of an acid salt lake: the role of biology in an extreme depositional environment. PLoS One 10, e0122869 (2015).

4 Zaikova, E., Benison, K. C., Mormile, M. R. & Johnson, S. S. Microbial communities and their predicted metabolic functions in a desiccating acid salt lake. Extremophiles 22, 367–379 (2018).

5 Futterer, O. et al. Genome sequence of *Picrophilus torridus* and its implications for life around pH 0. Proceedings of the National Academy of Sciences of the United States of America 101, 9091–9096 (2004).

6 Varet, J. in Geology of Afar {East Africa). Regional Geology Reviews (eds R. Oberhänsli, M. J. de Wit, & F. M. Roure) Ch. 7, 205–226 (Springer, 2018).

7 Franzson, H., Helgadóttir, H. M. & Óskarsson, F. in Proceedings World Geothermal Congress. 11.

8 Darrah, T. H. et al. Gas chemistry of the Dallol region of the Danakil Depression in the Afar region of the northern-most East African Rift. Chemical Geology 339, 16–29 (2013).

9 Holwerda, J. G. & Hutchinson, R. W. Potash-bearing evaporites in the Danakil area, Ethiopia. Economic Geology 63, 124–150 (1968).

10 Warren, J. K. Danakhil potash, Ethiopia: Beds of kainite/carnallite, Part 2 of 4. (2015).

11 Cavalazzi, B. et al. The Dallol geothermal area, Northern Afar (Ethiopia)-An exceptional planetary field analog on Earth. Astrobiology (2019).

12 Hallsworth, J. E. et al. Limits of life in MgCl2-containing environments: chaotropicity defines the window. Environ Microbiol 9, 801–813 (2007).

13 Stevenson, A. et al. Is there a common water-activity limit for the three domains of life? ISME J 9, 1333–1351 (2015).

14 McKay, C. P. Requirements and limits for life in the context of exoplanets. Proceedings of the National Academy of Sciences of the United States of America 111, 12628–12633 (2014).

15 Moissl-Eichinger, C., Cockell, C. & Rettberg, P. Venturing into new realms? Microorganisms in space. FEMS microbiology reviews 40, 722–737 (2016).

16 Pérez, E. & Chebude, Y. Chemical analysis of Gaet’ale, a hypersaline pond in Danakil Depression (Ethiopia): New record for the most saline water body on Earth. Aquat Geochem 23, 109–117 (2017).

17 Kotopoulou, E. et al. A polyextreme hydrothermal system controlled by iron: The case of Dallol at the Afar Triangle. ACS earth & space chemistry 3, 90–99 (2019).

18 Warren, J. K. Danakhil Potash, Ethiopia: Is the present geology the key? Part 1 of 4. (2015).

19 Tosca, N. J., Knoll, A. H. & McLennan, S. M. Water activity and the challenge for life on early Mars. Science (New York, N.Y.) 320, 1204–1207 (2008).

20 Stevenson, A. et al. Aspergillus penicillioides differentiation and cell division at 0.585 water activity. Environ Microbiol 19, 687–697 (2017).

21 Sheik, C. S. et al. Identification and removal of contaminant sequences from ribosomal gene databases: Lessons from the Census of Deep Life. Frontiers in microbiology 9 (2018).

22 Weyrich, L. S. et al. Laboratory contamination over time during low-biomass sample analysis. Molecular ecology resource (2019).

23 Narasingarao, P. et al. De novo metagenomic assembly reveals abundant novel major lineage of Archaea in hypersaline microbial communities. ISME J 6, 81–93 (2012).

24 Sorokin, D. Y. et al. Discovery of extremely halophilic, methyl-reducing euryarchaea provides insights into the evolutionary origin of methanogenesis. Nat Microbiol 2, 17081 (2017).

25 Sorokin, D. Y. et al. Methanonatronarchaeum thermophilum gen. nov., sp. nov. and ‘Candidatus Methanohalarchaeum thermophilum’, extremely halo(natrono)philic methyl-reducing methanogens from hypersaline lakes comprising a new euryarchaeal class Methanonatronarchaeia classis nov. International journal of systematic and evolutionary microbiology 68, 2199–2208 (2018).

26 Borin, S. et al. Sulfur cycling and methanogenesis primarily drive microbial colonization of the highly sulfidic Urania deep hypersaline basin. Proceedings of the National Academy of Sciences of the United States of America 106, 9151–9156 (2009).

27 Mwirichia, R. et al. Metabolic traits of an uncultured archaeal lineage--MSBL1--from brine pools of the Red Sea. Scientific reports 6, 19181 (2016).

28 Castelle, C. J. et al. Biosynthetic capacity, metabolic variety and unusual biology in the CPR and DPANN radiations. Nature reviews. Microbiology 16, 629–645 (2018).

29 Castelle, C. J. & Banfield, J. F. Major new microbial groups expand diversity and alter our understanding of the Tree of Life. Cell 172, 1181–1197 (2018).

30 Dombrowski, N., Lee, J. H., Williams, T. A., Offre, P. & Spang, A. Genomic diversity, lifestyles and evolutionary origins of DPANN archaea. FEMS Microbiol Lett 366(2019).

31 Petitjean, C., Deschamps, P., Lopez-Garcia, P. & Moreira, D. Rooting the domain archaea by phylogenomic analysis supports the foundation of the new kingdom proteoarchaeota. Genome biology and evolution 7, 191–204 (2014).

32 Golyshina, O. V. et al. ‘ARMAN’ archaea depend on association with euryarchaeal host in culture and in situ. Nature communications 8, 60 (2017).

33 Minegishi, H. et al. Acidophilic haloarchaeal strains are isolated from various solar salts. Saline systems 4, 16 (2008).

34 Naor, A. & Gophna, U. Cell fusion and hybrids in Archaea: prospects for genome shuffling and accelerated strain development for biotechnology. Bioengineered 4, 126–129 (2013).

35 Garcia-Ruiz, J. M. et al. Self-assembled silica-carbonate structures and detection of ancient microfossils. Science (New York, N.Y.) 302, 1194–1197 (2003).

36 Garcia-Ruiz, J. M., Melero-Garcia, E. & Hyde, S. T. Morphogenesis of self-assembled nanocrystalline materials of barium carbonate and silica. Science (New York, N. Y.) 323, 362–365 (2009).

37 Slonczewski, J. L., Fujisawa, M., Dopson, M. & Krulwich, T. A. Cytoplasmic pH measurement and homeostasis in bacteria and archaea. Advances in microbial physiology 55, 1–79, 317 (2009).

38 Buetti-Dinh, A., Dethlefsen, O., Friedman, R. & Dopson, M. Transcriptomic analysis reveals how a lack of potassium ions increases Sulfolobus acidocaldarius sensitivity to pH changes. Microbiology (Reading, England) 162, 1422–1434 (2016).

39 Gunde-Cimerman, N., Plemenitas, A. & Oren, A. Strategies of adaptation of microorganisms of the three domains of life to high salt concentrations. FEMS microbiology reviews 42, 353–375 (2018).

40 Lee, C. J. D. et al. NaCl-saturated brines are thermodynamically moderate, rather than extreme, microbial habitats. FEMS microbiology reviews 42, 672–693 (2018).

41 Colman, D. R., Lindsay, M. R. & Boyd, E. S. Mixing of meteoric and geothermal fluids supports hyperdiverse chemosynthetic hydrothermal communities. Nature communications 10, 681 (2019).

42 López-García, P., Zivanovic, Y., Deschamps, P. & Moreira, D. Bacterial gene import and mesophilic adaptation in archaea. Nature reviews. Microbiology 13, 447–456 (2015).

43 Hensel, R. & König, H. Thermoadaptation of methanogenic bacteria by intracellular ion concentration. FEMS Microbiology Letters 49, 75–79 (1988).

44 Shima, S., Thauer, R. K. & Ermler, U. Hyperthermophilic and salt-dependent formyltransferase from Methanopyrus kandleri. Biochemical Society transactions 32, 269–272 (2004).

45 Cray, J. A., Russell, J. T., Timson, D. J., Singhal, R. S. & Hallsworth, J. E. A universal measure of chaotropicity and kosmotropicity. Environ Microbiol 15, 287–296 (2013).

46 Cray, J. A. et al. Chaotropicity: a key factor in product tolerance of biofuel-producing microorganisms. Current opinion in biotechnology 33, 228–259 (2015).

47 Fox-Powell, M. G., Hallsworth, J. E., Cousins, C. R. & Cockell, C. S. Ionic strength is a barrier to the habitability of Mars. Astrobiology 16, 427–442 (2016).

48 R: A language and environment for statistical computing. v. http://www.r-project.org (R Foundation for Statistical Computing, Vienna, Austria, 2017).

49 Lê, S., Josse, J. & Husson, F. FactoMineR: An R package for multivariate analysis. Journal of Statistical Software 25, 1–18 (2008).

50 factoextra: Extract and visualize the results of multivariate data analyses (https://CRAN.R-project.org/package=factoextra, 2017).

51 Magoc, T. & Salzberg, S. L. FLASH: fast length adjustment of short reads to improve genome assemblies. Bioinformatics 27, 2957–2963 (2011).

52 Martin, M. Cutadapt removes adapter sequences from high-throughput sequencing reads. EMBnet.Journal 17, 10–12 (2011).

53 Rognes, T., Flouri, T., Nichols, B., Quince, C. & Mahe, F. VSEARCH: a versatile open source tool for metagenomics. PeerJ 4, e2584 (2016).

54 Fu, L., Niu, B., Zhu, Z., Wu, S. & Li, W. CD-HIT: accelerated for clustering the next-generation sequencing data. Bioinformatics 28, 3150–3152 (2012).

55 Quast, C. et al. The SILVA ribosomal RNA gene database project: improved data processing and web-based tools. Nucleic Acids Res 41, D590–596 (2013).

56 Guillou, L. et al. The Protist Ribosomal Reference database (PR2): a catalog of unicellular eukaryote Small Sub-Unit rRNA sequences with curated taxonomy. Nucleic Acids Res 41, D597–D604 (2013).

57 Katoh, K. & Standley, D. M. MAFFT multiple sequence alignment software version 7: improvements in performance and usability. Molecular biology and evolution 30, 772–780 (2013).

58 Capella-Gutierrez, S., Silla-Martinez, J. M. & Gabaldon, T. trimAl: a tool for automated alignment trimming in large-scale phylogenetic analyses. Bioinformatics 25, 1972–1973 (2009).

59 Nguyen, L. T., Schmidt, H. A., von Haeseler, A. & Minh, B. Q. IQ-TREE: a fast and effective stochastic algorithm for estimating maximum-likelihood phylogenies. Molecular biology and evolution 32, 268–274 (2015).

60 Asnicar, F., Weingart, G., Tickle, T. L., Huttenhower, C. & Segata, N. Compact graphical representation of phylogenetic data and metadata with GraPhlAn. PeerJ 3, e1029 (2015).

61 Wang, H. C., Xia, X. & Hickey, D. Thermal adaptation of the small subunit ribosomal RNA gene: a comparative study. Journal of molecular evolution 63, 120–126 (2006).

62 Rodriguez-Valera, F., Ruiz-Berraquero, F. & Ramos-Cormenzana, A. Behaviour of mixed populations of halophilic bacteria in continuous cultures. Canadian journal of microbiology 26, 1259–1263 (1980).

