## Supplementary Material for "Hyperdiverse archaea near life limits at the polyextreme geothermal Dallol area"

**This PDF file includes:**

Supplementary Figs. 1 to 6
Supplementary Tables 1 to 6

**Other Supplementary Materials for this manuscript include the following:**

Supplementary Data 1 to 7

**Supplementary Data 1 |** (Table) Organic and ionic chromatography analysis of samples from the Dallol dome and surrounding area.
**Supplementary Data 2 |** (Table) Identification, phylogenetic affinity and relative abundance of prokaryotic OTUs.
**Supplementary Data 3 |** Identification, phylogenetic affinity and relative abundance of eukaryotic OTUs from Dallol area samples.
**Supplementary Data 4 |** Full tree of archaeal 16S rRNA gene fragments in Newick format.
**Supplementary Data 5 |** Full tree of bacterial 16S rRNA gene fragments in Newick format.
**Supplementary Data 6 |** (Figures) Chemical maps obtained from energy dispersive X-ray spectrometry (EDXS) combined to SEM observations of cells and abiotic biomorphs observed in the Dallol area and Lake Assale (Karum).
**Supplementary Data 7 |** (Figures) Energy dispersive X-ray spectrometry (EDXS) spectra of different structures observed in samples from the Dallol area.

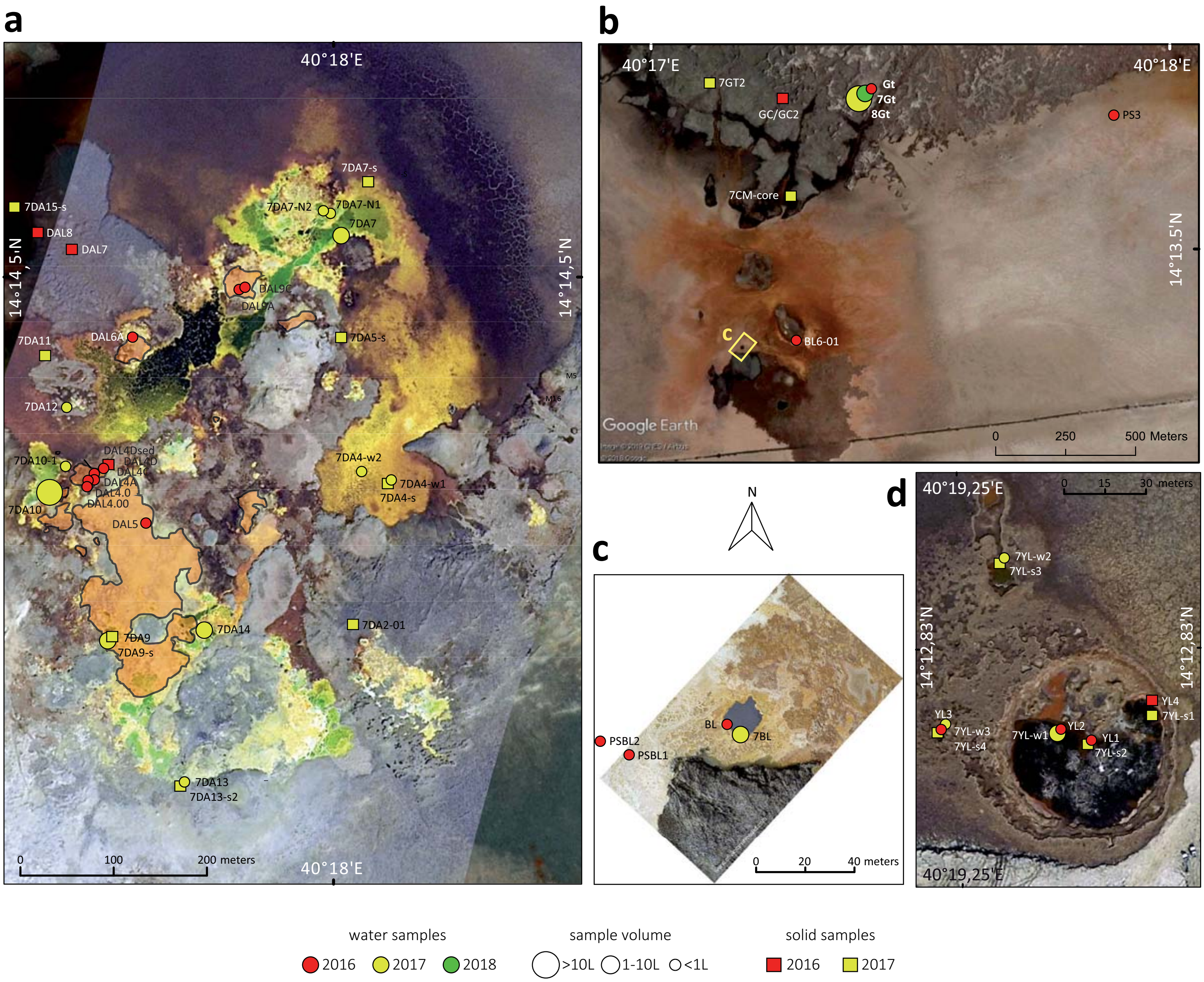

**Supplementary Fig. 1 | Aerial view of the main sampling sites in the Dallol area. a**, Dallol dome summit showing the acidic green-yellow-brown colored hydrothermal ponds and active degassing areas during our 2017 sampling trip; the orange-shaded area shows the active hydrothermal zone in January 2016. **b**, Dallol West salt canyons and Black Mountain area. **c**, Black Lake. **d**, Yellow Lake and surroundings. Names of samples and sampling sites are indicated. The size of circles is proportional to the water volume collected or filtered for subsequent analyses. Aerial photographs were taken from a drone by O. Grunewald, except b, which is a Google Earth aerial image (09/03/2016) provided by Image © 2019 CNES/Airbus.

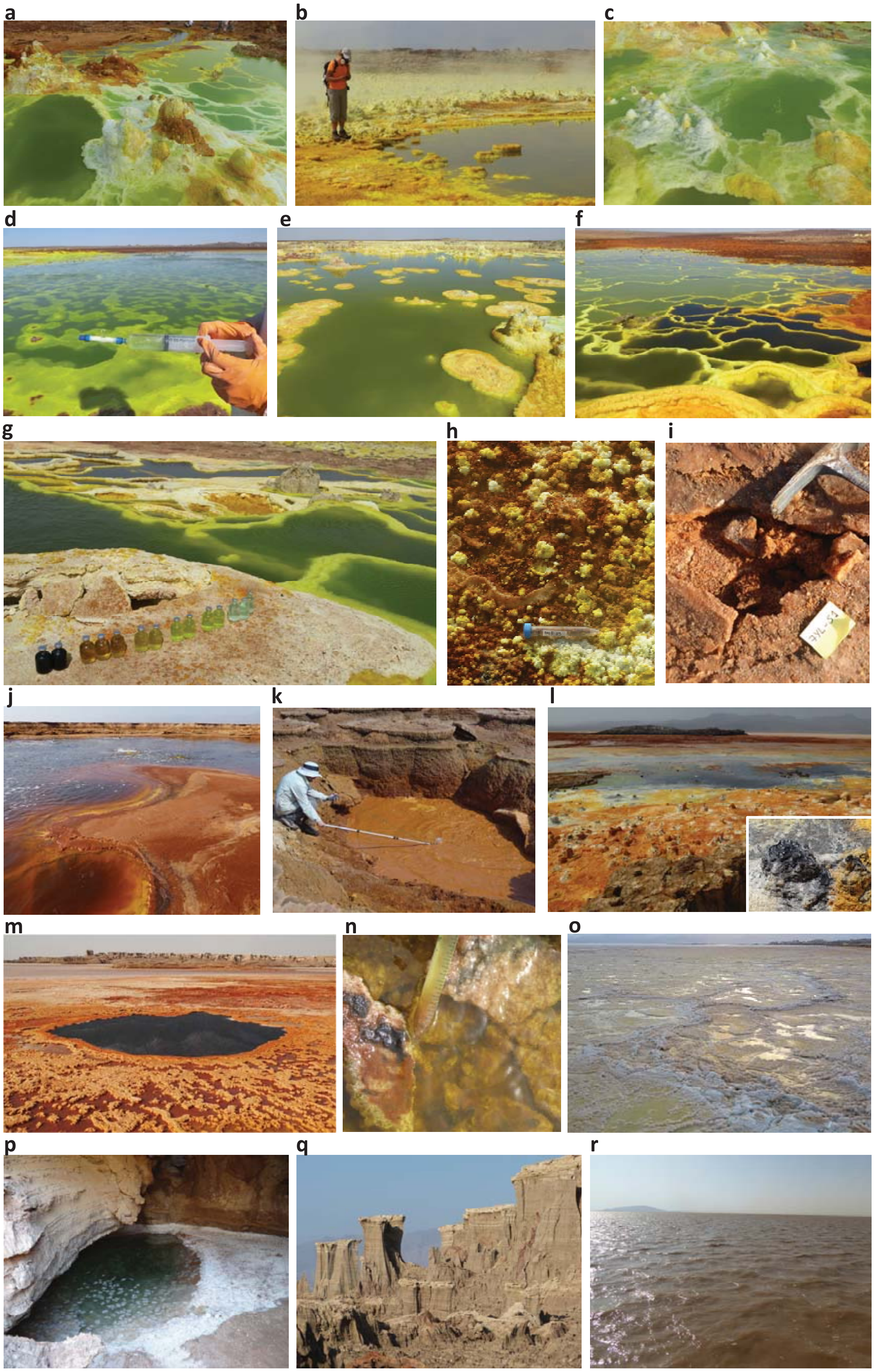

Supplementary Fig. 2 | Views of different sampling sites in the Dallol dome and surroundings in the Danakil Depression. **a**, DAL4 sampling site ponds; **b**, DAL5 pond and active degassing area; **c**, active hydrothermal springs in DAL9 ponds; **d**, in situ cell-trap filtration at the 7DA7 sampling area; **e**, 7DA9 sampling site; **f**,7DA10 ponds showing increasingly darker and brownish colors along the oxidation gradient; **g**, water samples from the different 7DA10 ponds; **h**, DAL8 mineral precipitates; **i**, 'proto-soil'-like salt crust (7YL-S1) near the Yellow Lake; **j**, Yellow Lake showing active degassing; **k**, YL3, salt-mud volcano in the Yellow Lake area; **l**, 'Little Dallol' hydrothermal very active area in 2016 on the way to the Black Mountain (in the distance; inlet, chimney emitting hydrocarbon-rich fluids at 110°C); **m**, Black Lake; **n**, PSBL2 (Black Lake area ponds); **o**, wet salt plain, influenced by hydrothermal activity, corresponding to PS3 sample area; **p**, the cave in the salt canyons where Gt, 7Gt and 8Gt samples were collected; **q**, salt canyons; **r**, Assale (Karum) lake. Sample names starting by 7 indicate collection in 2017. Pictures from all other samples/sampling sites were taken during the 2016 expedition.

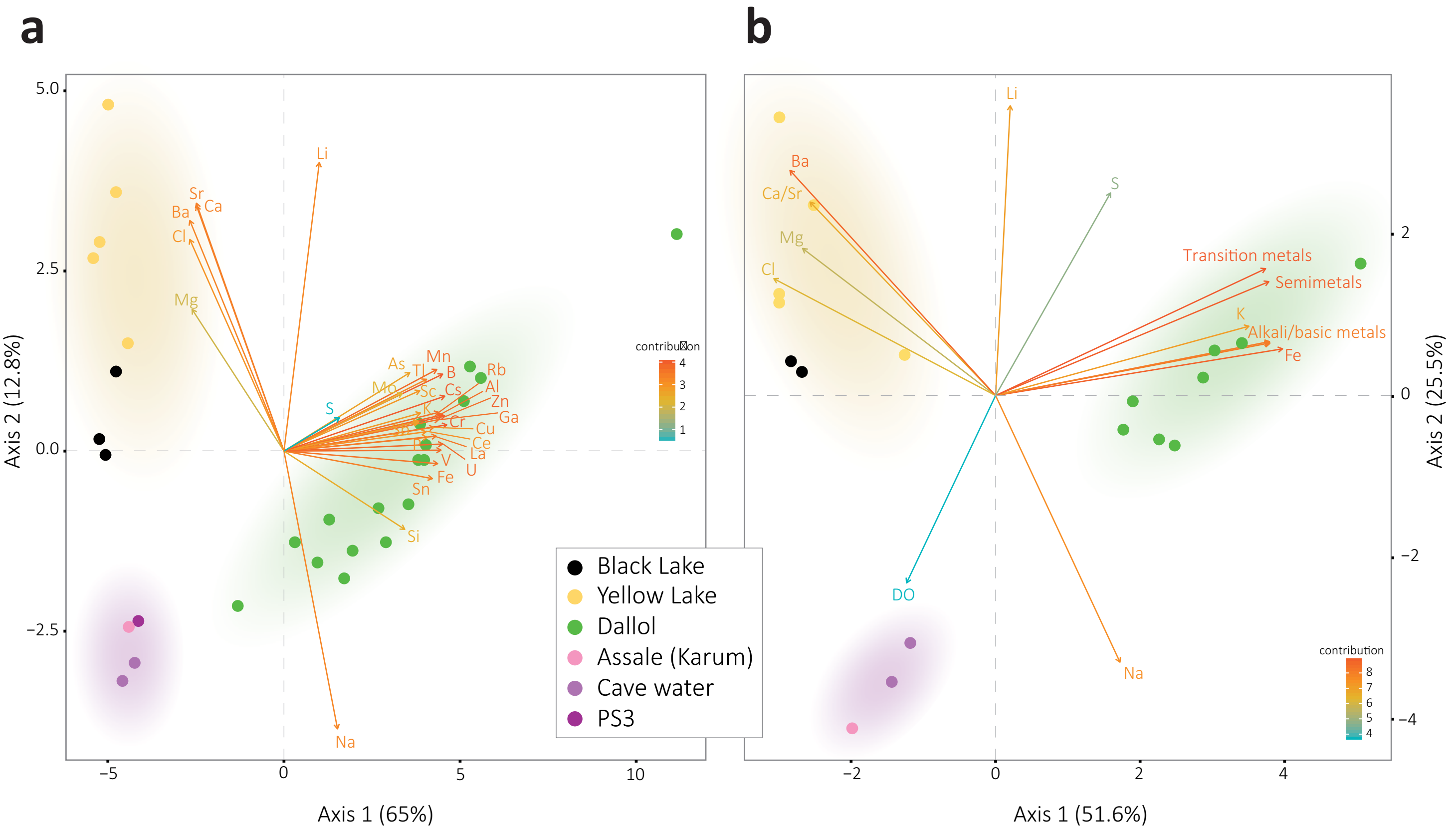

**Supplementary Fig. 3 | Principal Component Analyses (PCA) of Dallol area sampling sites as a function of physicochemical parameters.** PCA of 29 samples according to their chemical composition; only relatively abundant elements (see Supplementary Table 2) are included in the analysis. A summary of this analysis is shown in Fig.2f. **b**, PCA including the same variables as Fig. 2f but additionally including dissolved oxygen (DO). Measured parameters on site can be found in Supplementary Table 1. Colored zones in PCA analyses correspond to the three major chemical zones identified in this study.

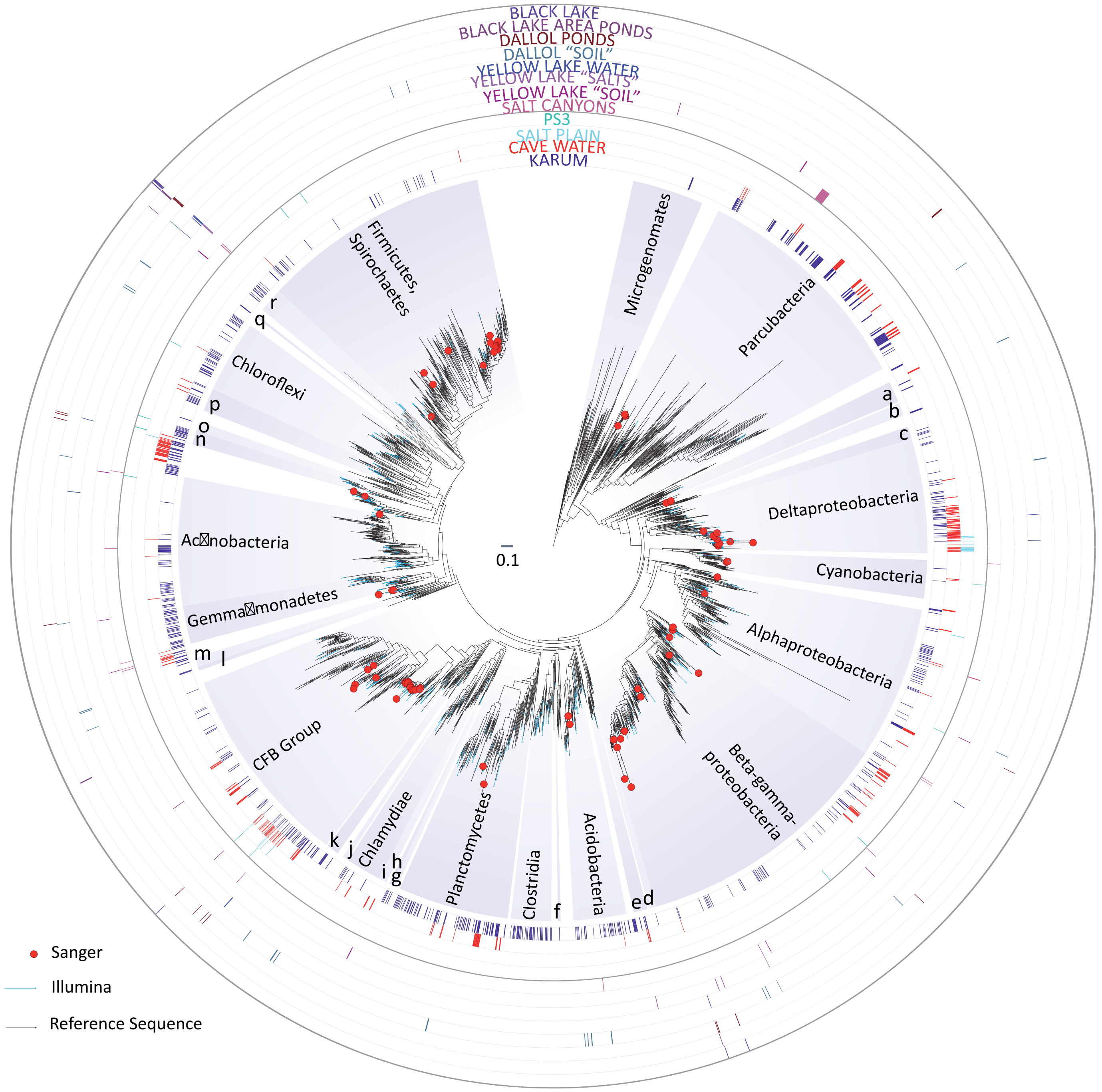

a. Peregrinibacteria
b. Saccharibacteria
c. Desulfovibrionales
d. Deferribacteres
e. Candidate division TM6
f. Candidate division NC10
g. Candidatus Latescibacteria

h. Candidatus BRC1
i. Omnitrophica
j. Zixibacteria
k. Marinimicrobia
l. Rokubacteria
m. Nitrospirae

n. Acetothermia
o. Synergistetes
p. Armatimonadetes
q. Dadabacteria
r. Deinococcus-Thermus

**Supplementary Fig. 4 | Phylogenetic tree of bacterial 16S rRNA gene sequences showing the phylogenetic placement of OTUs identified in the different Dallol area samples.** Sequences derived from metabarcoding studies are represented by blue lines (Illumina sequences); those derived from cloning and Sanger sequencing of environmental samples, cultures and FACS-sorted cells are labelled with a red dot. Reference sequences are in black. Concentric circles around the tree indicate the presence/absence of the corresponding OTUs in different groups of samples (groups shown in Fig.3a). Only sequences not deemed contaminant (see Supplementary Data 2-3) were included in the tree. The full tree is provided as Supplementary Data 5.

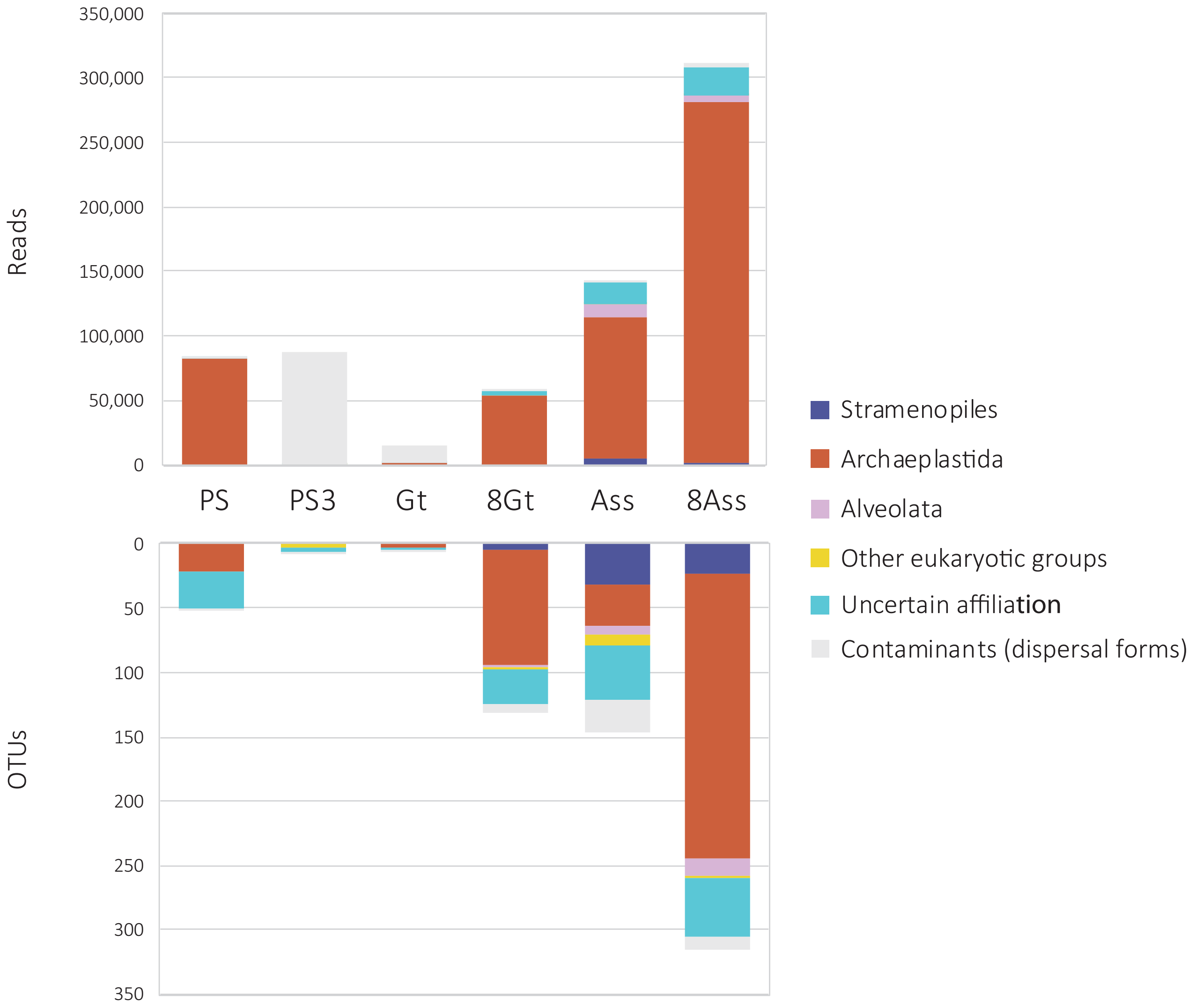

**Supplementary Fig. 5 | Eukaryotic presence, diversity and relative abundance in Dallol area samples.** Histogram showing the phylogenetic affiliation and abundance of 18S rRNA gene amplicon reads of eukaryotes (upper panel) obtained with universal eukarytic primers and the associated OTU diversity (lower panel). Only a few samples yielded amplicons; negative PCR controls were always negative. Sequences corresponding to macroscopic plants and fungi (probably derived from pollen or spores) were considered contaminant (light grey). The phylogenetic affiliation of dominant eukaryotic groups is color-coded.

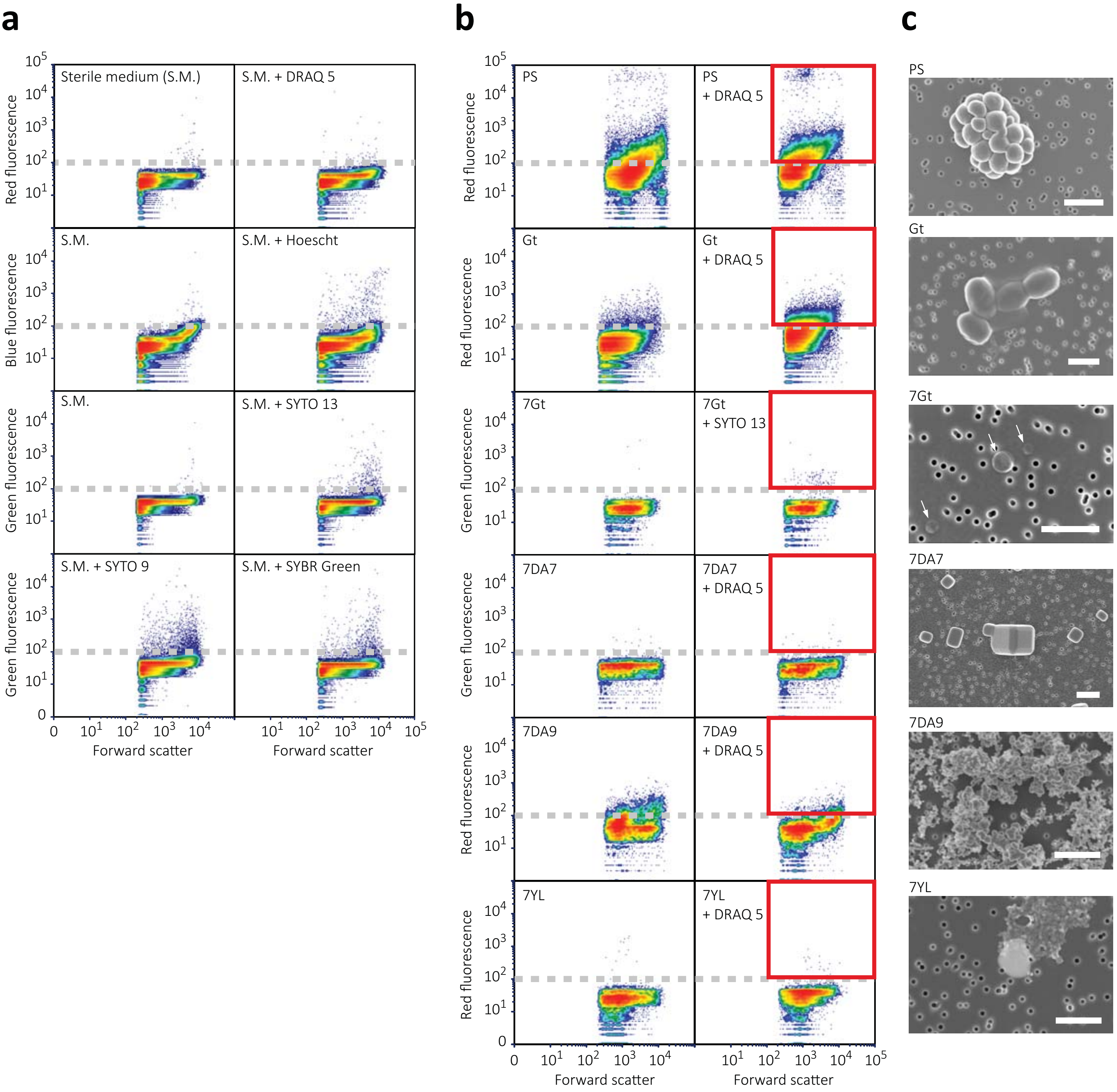

**Supplementary Fig. 6 | Multiparametric fluorescence analyses and fluorescence-activated cell sorting (FACS) analyses of representative Dallol area samples. a**, effect of DNA fluorescent dyes on background fluorescence emission; natural (left panels) and DNA dye-induced (right panels) fluorescence in the sterile hypersaline SALT-YE medium used to dilute/sort Dallol samples. Fluorescence is plotted against the size of the analyzed particles (forward scatter); events concentration is color-coded, red being high concentration and blue, low concentration. DRAQ 5 and SYTO 13 introduced less background and were chosen for FACS of natural samples. The approximate background threshold (ca. $10^2$) is indicated by a broken grey line. **b**, multiparametric fluorescence analyses of different Dallol samples before (left panels) and after (right panels) adding fluorescent DNA dyes. Events (particles) above background (red squares) were FACS-sorted and filtered on 0.1 µm pore-size filters prior to SEM observations. **c**, SEM photographs showing examples of sorted particles. Cells are observed in samples PS, Gt and 7Gt; halite crystals in 7DA7 and amorphous mineral particles in 7DA9 and 7YL. Arrows indicate ultrasmall cells. The scale bar is 1 µm.

Supplementary Table 1. **List and description of samples from the Dallol area analyzed in this study and type of analyses performed.** DO, dissolved oxygen; ORP, oxido-reduction potential; SEM/XRD, scanning electron microscopy/x-ray diffraction analyses; FACS, fluorescence-activated cell sorting analysis; n.a., not applicable; n.d. not determined. Refractometry-derived salinity refers to the percentage (w/v) of local salt composition (see Supplementary Table 2 and Supplementary Data 1 for elementary and ionic analyses) measured in situ. Salinity was also directly measured by weighting the total solids (dry weight experimentally measured in triplicates; SD, standard deviation).

| Color of ponds or solid samples | Sample name | Coordinates | Collection date | Brief description | Sample type / volume | | | Physicochemical parameters | | | | | | | Type of analyses | | | | | |
|---|---|---|---|---|---|---|---|---|---|---|---|---|---|---|---|---|---|---|---|---|
| | | | | | Solid | Liquid (ml) | 0.2-30 µm fraction (ml)[a] | Temp (°C) | pH | DO (%) | DO (mg/l) | ORP (mV) | Refractometry inferred salinity (%) | Salinity - Total Solids (g/l) ± SD | Chemistry | Cloning/ Sanger seq. | Meta-barcoding | Culture assays[b] | SEM/ XRD | FACS |
| **Dallol hydrothermal ponds** | | | | | | | | | | | | | | | | | | | | |
| | DAL4.00 | 14.23916N 40.297059E | 16.01.2016 | Hydrothermal fluid from a salt chimney feeding cascading ponds along redox gradient site DAL4 | | 50 | 1000 | 108.0 | n.d | n.d | n.d | n.d | 30.0 | 375.40 ± 10.06 | X | X | | | | |
| | DAL4.0 | 14.23923N 040.29707E | 17.01.2016 | Warm whitish green pond in the redox pond series DAL4 | | 50 | 1000 | 46.3 | -0.43 | 0.55 | 0.03 | 328.95 | 21.0 | 373.29 ± 4.83 | X | X | X | Y | X | |
| | DAL4A | 14.23921N 040.29713E | 17.01.2016 | Bright green pond in the redox pond series DAL4 | | 50 | 1000 | 30.5 | -0.53 | 3.05 | 0.26 | 366.8 | 22.0 | n.d | X | X | X | Y | X | |
| | DAL4C | 14.23967N 040.29744E | 16.01.2016 | Brownish green pond in the redox pond series DAL4 | | | 1000 | 31.6 | -0.95 | 1 | 0.08 | 384.05 | 21.5 | n.d | | X | X | Y | | |
| | DAL4D | 14.23934N 040.29728E | 16.01.2016 | Dark brown pond along an oxidation gradient, DAL4 site | | 50 | 1000 | 31.0 | -0.72 | 1.4 | 0.11 | 381.55 | n.d | 405.32 ± 19.6 | X | X | X | Y | X | |
| | DAL 4D-SED | 14.23934N 040.29728E | 16.01.2016 | Yellow salt front forming pond wall | X | | | n.a | n.a | n.a | n.a | n.a | n.a | n.a | X | X | | | | |
| | DAL 5 | 14.23869N 40.29776E | 17.01.2016 | Yellow water from one big Dallol pond | | 50 | | 42.6 | -0.93 | 3.4 | 0.19 | 373.3 | 40 | 386.84 ± 6.18 | | | | Y | | |
| | DAL6A | 14.24083N 040.29756E | 18.01.2016 | Fluid from a salt chimney feeding various ponds along an oxidation gradient, site DAL6 | | 50 | 1000 | 108.4 | -0.65 | n.d | n.d | n.d | n.d | 424.51 ± 38.22 | X | X | X | Y | | |
| | DAL 7 | 14.242343N 40.296798E | 18.01.2016 | Sulfur-colored mineral precipitates on "chocolate formation" | X | | | n.a | n.a | n.a | n.a | n.a | n.a | n.a | | | | Y | | |
| | DAL8-01 | 14.24217N 040.29635E | 20.01.2016 | Yellow, potentially sulfur-rich precipitates | X | | | n.a | n.a | n.a | n.a | n.a | n.a | n.a | | X | X | Y | | |
| | DAL8-02 | 14.24217N 040.29635E | 20.01.2016 | Golden, potentially sulfur-rich salt precipitates | X | | | n.a | n.a | n.a | n.a | n.a | n.a | n.a | | X | X | | | |
| | DAL8-03 | 14.24217N 040.29635E | 20.01.2016 | Brown, potentially sulfur-rich salt precipitates | X | | | n.a | n.a | n.a | n.a | n.a | n.a | n.a | | X | X | | | |
| | DAL9A | 14.241504N 40.298836E | 21.01.2016 | Large ligth green bubbling pond with various hydrothermal sources; site DAL9 | | 50 | 650 | 37.9 | -0.27 | n.d | n.d | n.d | n.d | n.d | X | X | X | Y | | |
| | DAL9C | 14.24151N 40.298874E | 21.01.2016 | Smaller deep green hydrothermal pond at the DAL9 site | | 50 | 750 | 56.0 | -0.20 | n.d | n.d | n.d | n.d | n.d | X | X | X | | | |
| | 7DA2-01 | 14.241275N 40.300633E | 06.01.2017 | Grey cauliflower-like mineral precipitates | X | | | n.a | n.a | n.a | n.a | n.a | n.a | n.a | | X | X | | | |
| | 7DA4-w1 | 14,239406N 40,300419E | 10.01.2017 | Yellow-brownish water from a drying pond formed by yellow-colored salt | | 100 | | 27.9 | -0.73 | 3.8 | 0.13 | 416.1 | 44 | 446.07 ± 11.41 | X | | | | | |
| | 7DA4-w2 | 14,239892N 40,300106E | 10.01.2017 | Slightly lighter water from the same kind of pool as 7DA4-w1 | | 100 | | 27.0 | -0.72 | 2.9 | 0.09 | 420.1 | 40 | 371.77 ± 2.11 | X | | | | | |
| | 7DA4-s | 14,239406N 40,300419E | 10.01.2017 | Fresh, yellow cauliflower-like mineral precipitates near 7DA4-w2 | X | | | n.a | n.a | n.a | n.a | n.a | n.a | n.a | | X | X | | | |
| | 7DA5-s | 14,240908N 40,300119E | 10.01.2017 | Brown (oxidized), fluid-impregnated cauliflower-like mineral precipitates | X | | | n.a | n.a | n.a | n.a | n.a | n.a | n.a | X | X | X | | | |
| | 7DA7 | 14.24219N 40.300102E | 07.01.2017 | Water from an active green large pond fed by multiple hydrothermal springs | | 500 | 3000 | 19.7 | -0.55 | 4.1 | 0.16 | 371.9 | 47 | 429.6 ± 12.89 | | X | X | X | X | X |
| | 7DA7-s | 14.2428N 40.300494E | 10.01.2017 | Brown salt crust East to 7DA7 active site on a drying pondcovering a layer of wet yellow salt | X | | | n.a | n.a | n.a | n.a | n.a | n.a | n.a | X | | | | | |
| | 7DA7N-1 | 14.242381N 40.299992E | 10.01.2017 | Bluish hot pond on the northern terrace feeding 7DA7 | | 50 | | 44.6 | -0.33 | 5.7 | 0.16 | 271.2 | 40.0 | 404.13 ± 20.16 | X | | | Y | | |
| | 7DA7N-2 | 14.242381N 40.299992E | 10.01.2017 | Hotter, greenish pond adjacent to 7DA7N-1 | | 50 | | 68.0 | n.d | n.d | n.d | n.d | 37.0 | 411.1 ± 0.57 | X | | | | | |
| | 7DA9 | 14.241528N 40.29884E | 08.01.2017 | Large pond in active degassing area with many white-yellowish chimneys and salt nenuphars | | 500 | 6000 | 31.9 | -0.34 | 5.5 | 0.16 | 369 | 43 | 379.03 ± 17.18 | X | X | X | X | X | X |
| | 7DA9-s | 14.241528N 40.29884E | 08.01.2017 | Centimetric round-shaped salt and sulfur formations in a dry pond next to 7DA9 | X | | | n.a | n.a | n.a | n.a | n.a | n.a | n.a | | X | X | | | |
| | 7DA10 | 14.23908N 40.296598E | 09.01.2017 | Upper green pond in a well-marked terrace system along an oxidation gradient | | 100 | 14700 | 55.2 | -0.08 | 2.2 | 0.05 | 321.8 | 35.0 | 366.83 ± 7.71 | X | X | X | | | |
| | 7DA10-1 | 14.23908N 40.296598E | 09.01.2017 | Highly evaporated, dark brown pond, lower in the 7DA10 terrace system | | 100 | | 33.4 | n.d | n.d | n.d | n.d | 70 | n.d | X | | | Y | | |
| | 7DA12 | 14.240136N 40.29675E | 10.01.2017 | Active chimney (nearby 2016 site DAL6) | | 100 | | 108.3 | 0.47 | n.d | n.d | n.d | 43.0 | 385.48 ± 21.18 | X | | | | | |
| | 7DA13-w1 | 14.235495N 40.298135E | 11.01.2017 | Hydrothermal fluid from big grey active chimney | | 100 | | 103.6 | n.d | n.d | n.d | n.d | 35.0 | 342.50 ± 18.87 | X | | | | | |
| | 7DA13-s2 | 14.235495N 40.298135E | 11.01.2017 | Grey, hard salt fragments at the bottom of chimney 7DA13 | X | | | n.a | n.a | n.a | n.a | n.a | n.a | n.a | | X | X | | | |
| | 7DA14 | 14.237398N 40.298428E | 12.01.2017 | Central pond in active group of geothermal ponds. Green color | | 50 | 675 | 39.7 | -0.51 | 2.4 | 0.15 | 382 | 40 | 338.90 ± 8.17 | X | X | X | | | |
| | 7DA15-s | 14.2421427N 40.297727E | 13.01.2017 | Green/yellow sediment near the "chocolate formation" CH-F | X | | | n.a | n.a | n.a | n.a | n.a | n.a | n.a | | X | X | | | |
| **Yellow Lake (Gaet'Ale)** | | | | | | | | | | | | | | | | | | | | |
| | YL1 | 14.213574N 40.32128E | 19.01.2016 | Yellow Lake bubbling water, yellow-orange color, oily textyre, smell of organics-containing gas | X | 50 | 400 | 40.6 | 1.88 | 6.6 | 0.37 | 447.1 | >50 | 724.40 ± 5.77 | X | X | X | Y | X | |
| | YL2 | 14.213648N 40.321188E | 19.01.2016 | Yellow Lake bubbling water - nearby spot (few meters away) | | 50 | 400 | 39.5 | 1.88 | 5.4 | 0.35 | 419.3 | 51.0 | 677.60 ± 2.24 | X | X | X | Y | | |
| | YL3-01-w | 14.213598N 40.320748E | 19.01.2016 | Water from salt/sediment volcano | | | 400 | 37.1 | n.d | n.d | n.d | n.d | >50 | 761.59 ± 69.51 | | X | X | Y | X | |
| | YL3-01-s | 14.213598N 40.320748E | 19.01.2016 | Salty deposits from bubbling salt/sediment volcano - reddish color | X | | | 37.1 | n.a | n.a | n.a | n.a | n.a | n.a | | X | X | | | |
| | YL4-01 | 14.21386N 40.321482E | 19.01.2016 | Salt crust on top of sediment at the further upper rim of Yellow Lake | X | | | n.a | n.a | n.a | n.a | n.a | n.a | n.a | | X | X | | | |
| | 7YL-w1 | 14.213574N 40.32128E | 12.01.2017 | Water from the Yellow Lake | | 150 | 5000 | 37.4 | 1.52 | 11.5 | 0.63 | 462.2 | >50 | 944.60 ± 60.38 | X | | | X | X | |
| | 7YL-w2 | 14.213648N 40.321188E | 12.01.2017 | Water from a mid-sized pond near the Yellow Lake, strong smell of organics, dead birds | | 200 | | 33.8 | 2.40 | 4.4 | 0.20 | 310.3 | 70 | 683.80 ± 80.61 | X | | | X | | X |
| | 7YL-w3 | 14.213598N 40.320748E | 12.01.2017 | Water from salt mud volcano (YL3 in 2016) | | 100 | | 35.9 | 1.37 | 5.1 | 0.32 | 349.3 | 80 | n.d | X | | | X | | |
| | 7YL-s1 | 14.213574N 40.32128E | 12.01.2017 | Salt crust from dried area in Yellow Lake | X | | | n.a | n.a | n.a | n.a | n.a | n.a | n.a | | X | X | | | |
| | 7YL-s2 | 14.213648N 40.321188E | 12.01.2017 | Pinkish salt forming ripple marks on the rim of the Yellow Lake | X | | | n.a | n.a | n.a | n.a | n.a | n.a | n.a | | X | X | | | |
| | 7YL-s3 | 14.213598N 40.320748E | 12.01.2017 | Salt fragments from rim of 7YL-w2 | X | | | n.a | n.a | n.a | n.a | n.a | n.a | n.a | X | X | X | | | |
| | 7YL-s4 | 14.2136083N 40.3212611E | 12.01.2017 | Reddish salt/sediment from salt mud volcano | X | | | n.a | n.a | n.a | n.a | n.a | n.a | n.a | | X | X | | | |
| **Black Lake and surroundings** | | | | | | | | | | | | | | | | | | | | |
| | PSBL1 | 14.221732N 40.28582E | 19.01.2016 | Reddish salt and water from pond close to Black Lake | | 50 | | 40.6 | 2.63 | n.a | n.a | n.a | 62.0 | 643.40 ± 22.20 | | X | X | XY | | |
| | PSBL2 | 14.221788N 40.285713E | 19.01.2016 | Yellowish salt and water from pond close to Black Lake | | 50 | | 40.6 | 2.50 | n.a | n.a | n.a | 58.0 | 620.4 | | X | X | XY | | |
| | PSBL3 | 14.22173N 040.28582E | 19.01.2016 | Orange salt and water from pond between Black Lake and camp site | | 50 | | n.d | n.d. | n.a | n.a | n.a | 34.0 | 344.40 ± 10.77 | | | | XY | X | |
| | PSBL4 | 14.22173N 040.28582E | 22.01.2016 | Warm red pond with acid emissions | | 50 | | 40.0 | 3.50 | n.a | n.a | n.a | 38.0 | 366.67 ± 9.63 | | | | X | X | |
| | BL2 | 14.221842N 40.286173 E | 19.01.2016 | Black Lake bischofite-enriched water, very high viscosity | | 25 | | n.a | n.a | n.a | n.a | n.a | n.a | n.d | | X | | | | |
| | BL / BL1-02 / BL-w | 14.22182N 40.286208E | 19.01.2016 | Black Lake bischofite-enriched water, very high viscosity | | 50 | 150 | 70.6 | 3.50 | n.a | n.a | n.a | n.a | 819.83 ± 181.88 | X | X | X | Y | | |
| | BL6-01 | 14.22211N 40.28805E | 24.01.2016 | Black fluid from chimney in very active young geothermal formation, black dome area | | 15 | | 110.0 | 4.40 | | | | n.d | n.d | | | | Y | | |
| | 7BL-w1 | 14.22182N 40.286208E | 10.01.2017 | Black Lake bischofite-enriched water from surface | | 150 | | 60.1 | 2.57 | 2.8 | 0.11 | 226.9 | 76 | 718.94 ± 17.94 | X | | X | X | | |
| | 7BL-w2 | 14.22182N 40.286208E | 10.01.2017 | Black Lake bischofite-enriched water, 3 m depth | | 100 | | n.a | n.a | n.a | n.a | n.a | n.a | n.d | X | | X | Y | | |
| **Salt plain at Dallol dome base** | | | | | | | | | | | | | | | | | | | | |
| | PS | 14.2250194N 40.2889000E | 19.01.2016 | Salt pan fragment between the Dallol dome and Black Lake, rehydrated with sterile spring water | X | 100 | 30 | n.a | n.a | n.a | n.a | n.a | n.a | n.a | | X | X | X | X | X |
| | PS3 | 14.22912N 40.297935E | 19.01.2016 | Water from salt shallow, hydrothermally influenced, pond at the dome base | | 150 | 950 | 29.3 | 4.21 | n.d | n.d | n.d | 20.0 | 352.64 ± 10.78 | X | X | X | XY | X | |
| **Lake Assale (Lake Karum)** | | | | | | | | | | | | | | | | | | | | |
| | Ass | 14.089567N 40.348583E | 23.01.2016 | Water from Lake Assale (Karum), overflooded towards the North of the Danakil Depression | | | 8000 | 26.2 | 6.68 | 53.0 | 4.30 | 97.7 | 20.0 | 360.72 ± 11.52 | | X | X | X | | |
| | 8Ass | 14.089567N 40.348583E | 15.01.2018 | Water from Lake Assale (Lake Karum) | | 100 | 3000 | 22.0 | 6.54 | 14 | 2.82 | 221 | 31 | 363.15 ± 11.51 | X | X | | XY | X | |
| **Dallol Salt Canyons** | | | | | | | | | | | | | | | | | | | | |
| | Gt | 14.229704N 40.289952E | 18.01.2016 | Cave hypersaline water reservoir at the salt canyons, West Dallol dome | | 50 | 50 | n.d | n.d | n.d | n.d | n.d | 30 | n.d | | X | X | XY | X | X |
| | 7Gt | 14.229704N 40.289952E | 11.01.2017 | Cave hypersaline water reservoir at the salt canyons, West Dallol dome | | 100 | 10000 | 29.0 | 6.27 | 9.5 | 0.27 | 117.2 | 30 | 323.87 ± 19.39 | | X | X | XY | X | X |
| | 7Gt2 | 14.23043N 40.285295E | 13.01.2017 | Small core at the mouth of a dry subterranean stream bed at the Western Dallol salt canyons | X | | | n.a | n.a | n.a | n.a | n.a | n.a | n.d | | X | X | | | |
| | 8Gt | 14.23043N 40.285295E | 16.01.2018 | Cave hypersaline water reservoir at the salt canyons, West Dallol dome | | 100 | 3000 | 22.0 | 6.54 | 15.2 | 2.30 | 262.1 | 30 | 330.82 ± 10.78 | X | X | | X | X | |
| | GC/GC2 | 14.229658N 40.287723E | 15.01.2016 | Gypsum crust from Dallol West salt canyons | X | | | n.a | n.a | n.a | n.a | n.a | n.a | n.a | X | X | X | | | |
| | 7CM-core | 14.226668N 40.287813E | 13.01.2017 | Small core of hydrothermally-influenced orange salty mud core at the base od the dome | X | | | n.a | n.a | n.a | n.a | n.a | n.a | n.a | | X | X | | | |

[a] Volume of water filtered to retain particles/cells in the indicated size range

[b] Samples labeled X and Y represent independent culture assays in two different laboratories (X-Orsay, Y-Madrid)

**Supplementary Table 2. Chemical composition of samples from the multi-extreme area of Dallol.** For details about chemical analyses, see Methods. Data about organic compounds identified in Dallol samples are listed in Supplementary Data 1.

**Liquid samples**

Concentration (mg/l)

| Element | DAL4.00 | DAL4.0 | DAL4A | DAL4D | DAL6A | DAL9A | DAL9C | 7DAL4-W1 | 7DAL4-W2 | 7DAL7 | 7DAL7-N1 | 7DAL7-N2 | 7DAL9 | 7DAL10 | 7DAL12 |
|---|---|---|---|---|---|---|---|---|---|---|---|---|---|---|---|
| Cl | 105190 | 111510 | 22610 | 99430 | 126750 | 123950 | 136920 | 170680 | 144790 | 201890 | 192280 | 179610 | 149770 | 170150 | 191470 |
| Na | 93210.36 | 105537.04 | 85029.17 | 81641.76 | 107204.75 | 95246.84 | 114638.67 | 39699.19 | 54954.80 | 51558.63 | 56742.29 | 59563.51 | 64186.60 | 69668.23 | 87876.56 |
| Ca | 2263.99 | 2867.06 | 2788.01 | 3388.86 | 1055.45 | 2390.61 | 3059.76 | 7937.79 | 6169.13 | 4301.43 | 4767.66 | 3892.18 | 6084.12 | 3774.66 | 1158.84 |
| Mg | 3764.86 | 4550.75 | 4387.41 | 6127.92 | 4538.48 | 6269.98 | 6268.84 | 9790.36 | 6348.22 | 6440.81 | 7206.49 | 5872.36 | 4249.59 | 3099.97 | 3131.56 |
| Fe | 26445.16 | 33578.06 | 32990.09 | 44030.37 | 32940.60 | 41621.88 | 40293.36 | 46182.85 | 29064.79 | 45206.84 | 47342.72 | 39815.45 | 23675.95 | 23107.07 | 26107.32 |
| K | 9047.00 | 17096.11 | 16445.10 | 22355.01 | 16684.14 | 22301.91 | 21746.24 | 28073.69 | 17451.30 | 23683.71 | 24945.93 | 21539.29 | 13469.45 | 12000.32 | 12859.23 |
| S | 2027 | 2077 | 2529 | 2984 | 3479 | 3111 | 3070 | 1836 | 2306 | 1240 | 1180 | 881 | 1560 | 1112 | 1840 |
| Mn | 673.24 | 853.16 | 847.55 | 1141.84 | 858.47 | 1121.10 | 1129.63 | 1551.99 | 1023.92 | 1281.68 | 1343.48 | 1169.53 | 773.03 | 696.02 | 691.42 |
| Sr | 68.06 | 85.09 | 84.34 | 108.12 | 41.38 | 72.39 | 81.61 | 188.87 | 124.26 | 104.78 | 117.33 | 98.58 | 122.28 | 93.31 | 40.41 |
| B | 353.15 | 456.40 | 457.15 | 608.86 | 369.37 | 470.06 | 504.49 | 899.60 | 591.48 | 529.14 | 607.12 | 486.72 | 491.64 | 379.03 | 297.55 |
| Al | 280.02 | 355.55 | 357.62 | 460.10 | 190.25 | 356.38 | 387.96 | 1113.20 | 722.04 | 544.99 | 614.45 | 516.50 | 689.24 | 354.23 | 132.78 |
| V | 21.71 | 26.94 | 27.36 | 37.71 | 27.23 | 35.55 | 35.95 | 45.56 | 28.15 | 46.14 | 47.10 | 40.62 | 25.00 | 20.86 | 21.93 |
| Si | 30.58 | 28.07 | 14.89 | 20.88 | 21.31 | 23.81 | 30.95 | 23.55 | 13.79 | 21.22 | 23.17 | 17.47 | 24.29 | 12.20 | 23.16 |
| P | 16.50 | 27.10 | 29.26 | 33.31 | 14.99 | 14.65 | 14.31 | 52.04 | 23.41 | 56.09 | 40.18 | 28.93 | 14.28 | 26.25 | 24.02 |
| Li | 10.92 | 14.88 | 15.07 | 21.52 | 14.54 | 18.62 | 18.99 | 23.61 | 15.08 | 19.26 | 22.51 | 16.66 | 10.42 | 10.24 | 12.84 |
| Cu | 31.27 | 24.94 | 7.53 | 17.89 | 15.90 | 19.80 | 33.21 | 50.64 | 28.91 | 41.85 | 30.21 | 31.27 | 42.91 | 19.86 | 11.51 |
| Zn | 52.21 | 68.71 | 67.28 | 91.20 | 60.50 | 82.24 | 87.23 | 140.51 | 86.32 | 105.22 | 105.98 | 88.93 | 77.46 | 57.62 | 49.60 |
| Kr | 0.00 | 0.00 | 0.00 | 0.00 | 0.00 | 0.00 | 0.00 | 0.00 | 5.82 | 0.00 | 17.09 | 13.01 | 0.00 | 19.95 | 19.58 |
| Rb | 14.74 | 17.33 | 16.36 | 22.00 | 15.57 | 21.92 | 22.42 | 42.29 | 26.14 | 30.13 | 30.78 | 27.10 | 19.45 | 15.43 | 13.56 |
| Ba | 5.84 | 5.88 | 3.27 | 5.05 | 2.61 | 2.77 | 7.55 | 3.00 | 3.05 | 6.77 | 9.71 | 7.26 | 11.94 | 4.49 | 3.25 |
| Ti | 1.73 | 1.46 | 1.86 | 2.41 | 0.76 | 1.05 | 1.37 | 6.17 | 4.43 | 1.93 | 1.43 | 1.24 | 2.13 | 1.39 | 0.46 |
| Au | 3.90 | 34.93 | 12.83 | 7.89 | 4.56 | 3.01 | 2.04 | 0.00 | 0.00 | 0.00 | 0.00 | 0.00 | 0.00 | 0.00 | 0.00 |
| Ni | 0.10 | 10.48 | 0.20 | 11.60 | 0.02 | 11.14 | 12.20 | 0.53 | 0.34 | 0.35 | 0.51 | 0.39 | 11.29 | 10.95 | 0.00 |
| Cr | 0.92 | 1.19 | 1.14 | 1.37 | 0.69 | 1.12 | 1.37 | 3.05 | 1.98 | 1.77 | 1.84 | 1.62 | 1.78 | 1.11 | 0.69 |
| Se | 0.00 | 0.44 | 0.00 | 1.30 | 0.07 | 0.73 | 0.11 | 0.20 | 1.66 | 0.02 | 0.68 | 0.20 | 0.41 | 0.00 | 0.33 |
| Co | 0.60 | 0.73 | 0.77 | 1.15 | 0.19 | 0.90 | 1.00 | 1.89 | 1.22 | 2.08 | 2.51 | 2.13 | 0.93 | 0.50 | 0.06 |
| Sn | 0.90 | 0.80 | 0.31 | 0.51 | 0.47 | 0.44 | 0.69 | 0.84 | 0.48 | 0.61 | 0.51 | 0.46 | 0.63 | 0.48 | 0.37 |
| Ga | 0.34 | 0.57 | 0.43 | 0.61 | 0.34 | 0.43 | 0.55 | 1.13 | 0.64 | 0.61 | 0.75 | 0.69 | 0.21 | 0.05 | 0.32 |
| Pb | 0.07 | 0.21 | 0.17 | 0.00 | 0.46 | 0.29 | 0.33 | 0.01 | 0.00 | 0.00 | 0.33 | 1.15 | 0.06 | 0.31 | 1.33 |
| As | 0.20 | 0.15 | 0.08 | 0.15 | 0.28 | 0.15 | 0.07 | 1.84 | 0.86 | 0.83 | 0.42 | 0.51 | 0.49 | 0.38 | 0.75 |
| Cs | 0.26 | 0.31 | 0.32 | 0.42 | 0.28 | 0.39 | 0.41 | 0.82 | 0.53 | 0.53 | 0.56 | 0.46 | 0.37 | 0.27 | 0.23 |
| U | 0.25 | 0.34 | 0.33 | 0.46 | 0.34 | 0.38 | 0.38 | 0.65 | 0.39 | 0.58 | 0.57 | 0.52 | 0.58 | 0.37 | 0.36 |
| Ce | 0.34 | 0.42 | 0.39 | 0.41 | 0.10 | 0.24 | 0.34 | 0.77 | 0.59 | 0.36 | 0.30 | 0.25 | 0.53 | 0.45 | 0.09 |
| La | 0.35 | 0.45 | 0.39 | 0.48 | 0.12 | 0.26 | 0.39 | 0.75 | 0.53 | 0.40 | 0.37 | 0.31 | 0.51 | 0.47 | 0.11 |
| Xe | 0.00 | 0.00 | 0.00 | 0.00 | 0.00 | 0.00 | 0.00 | 0.00 | 0.00 | 0.79 | 0.00 | 0.00 | 0.00 | 0.00 | 0.00 |
| Sc | 0.04 | 0.16 | 0.20 | 0.22 | 0.04 | 0.13 | 0.13 | 0.76 | 0.47 | 0.19 | 0.19 | 0.11 | 0.43 | 0.23 | 0.13 |
| Th | 0.01 | 0.56 | 0.23 | 0.17 | 0.07 | 0.08 | 0.06 | 0.11 | 0.10 | 0.20 | 0.00 | 0.01 | 0.58 | 0.16 | 0.00 |
| Mo | 0.11 | 0.03 | 0.01 | 0.03 | 0.00 | 0.00 | 0.00 | 0.37 | 0.28 | 0.32 | 0.28 | 0.28 | 0.15 | 0.07 | 0.03 |
| Sb | 0.18 | 0.12 | 0.02 | 0.07 | 0.06 | 0.07 | 0.09 | 0.36 | 0.21 | 0.08 | 0.06 | 0.09 | 0.21 | 0.11 | 0.08 |
| Tl | 0.06 | 0.08 | 0.09 | 0.11 | 0.07 | 0.10 | 0.10 | 0.17 | 0.11 | 0.10 | 0.12 | 0.09 | 0.10 | 0.06 | 0.04 |
| Be | 0.07 | 0.06 | 0.07 | 0.11 | 0.10 | 0.06 | 0.09 | 0.09 | 0.13 | 0.10 | 0.07 | 0.04 | 0.13 | 0.02 | 0.02 |

| Element | 7DAL13-W1 | 7DAL14 | BL | 7BL-W1 | 7BL-W2 | YL1 | YL1-2 (YL2) | 7YL-W1 | 7YL-W2 | 7YL-W3 | PS3 | 7Gt | 8GT | 8Ass |
|---|---|---|---|---|---|---|---|---|---|---|---|---|---|---|
| Cl | 161980 | 172310 | 194550 | 208200 | 220160 | 285390 | 373800 | 283480 | 255200 | 325980 | 149280 | 146110 | 230449 | 248886 |
| Na | 85356.59 | 61908.40 | 1764.03 | 1541.54 | 1607.32 | 2616.01 | 2375.56 | 1441.97 | 5149.17 | 1652.79 | 93924.71 | 89081.01 | 144416.34 | 112833.19 |
| Ca | 6123.51 | 6707.38 | 13436.28 | 8071.31 | 8306.92 | 163132.50 | 192095.04 | 175942.77 | 111085.94 | 162942.51 | 20316.50 | 6929.85 | 4037.16 | 23148.69 |
| Mg | 2123.95 | 5260.42 | 145295.05 | 108652.70 | 112683.58 | 53922.80 | 60599.66 | 41427.89 | 37276.85 | 42199.86 | 10953.71 | 1914.20 | 1778.91 | 4148.39 |
| Fe | 5337.40 | 24115.73 | 2719.04 | 1505.35 | 1488.73 | 91.65 | 74.52 | 81.98 | 418.42 | 324.10 | 0.00 | 2.18 | | 0.09 |
| K | 4421.73 | 14975.34 | 820.22 | 2323.18 | 2213.50 | 6186.46 | 4735.97 | 3834.52 | 24950.94 | 4617.63 | 7047.53 | 4737.88 | 2729.60 | 4759.99 |
| S | 2340 | 1543 | 1648 | ND | ND | 2427 | 3092 | 660 | 410 | 450 | 1384 | 351 | | |
| Mn | 283.56 | 860.66 | 393.63 | 206.67 | 208.93 | 381.84 | 442.58 | 362.36 | 251.53 | 385.35 | 89.69 | 1.54 | 1.61 | 5.11 |
| Sr | 77.33 | 118.49 | 331.62 | 186.03 | 186.68 | 3042.29 | 3462.02 | 2885.69 | 2032.27 | 2747.67 | 439.81 | 159.94 | 98.52 | 372.42 |
| B | 233.89 | 547.38 | 146.22 | 125.20 | 98.88 | 190.02 | 215.02 | 179.76 | 148.99 | 187.95 | 47.31 | 28.48 | 14.79 | 48.57 |
| Al | 297.94 | 641.44 | 0.71 | 0.00 | 0.00 | 0.72 | 0.89 | 0.00 | 0.00 | 0.00 | 1.32 | 0.00 | 0.03 | |
| V | 6.08 | 25.82 | 0.15 | 0.10 | 0.08 | 0.05 | 0.06 | 0.03 | 0.02 | 0.05 | 0.01 | 0.02 | 0.05 | 0.08 |
| Si | 28.98 | 25.02 | 13.85 | 4.30 | 2.31 | 5.65 | 8.46 | 9.62 | 0.00 | 8.15 | 9.30 | 10.87 | 3.00 | 11.12 |
| P | 4.07 | 11.22 | 0.00 | 3.01 | 2.05 | 0.00 | 0.00 | 2.45 | 0.76 | 1.11 | 0.00 | 1.56 | 6.88 | 7.81 |
| Li | 4.10 | 11.66 | 15.81 | 6.93 | 8.24 | 29.13 | 33.94 | 23.99 | 15.18 | 20.27 | 3.53 | 4.02 | 3.23 | 4.67 |
| Cu | 2.63 | 51.25 | 0.88 | 0.00 | 0.00 | 1.05 | 2.72 | 0.65 | 0.00 | 0.01 | 4.54 | 0.00 | 0.02 | 0.00 |
| Zn | 35.56 | 79.77 | 33.77 | 22.67 | 21.01 | 14.07 | 13.73 | 14.90 | 14.19 | 14.77 | 6.47 | 10.94 | 9.14 | 4.15 |
| Kr | 0.00 | 20.09 | 4.39 | 0.00 | 17.17 | 0.00 | 19.20 | 0.00 | 0.00 | 27.20 | 0.00 | 0.00 | | |
| Rb | 8.33 | 21.92 | 0.29 | 1.64 | 1.66 | 6.24 | 3.85 | 3.40 | 20.05 | 2.54 | 5.93 | 6.16 | 3.32 | 6.97 |
| Ba | 6.33 | 12.64 | 35.36 | 25.23 | 26.06 | 26.46 | 31.44 | 26.46 | 17.01 | 24.50 | 7.67 | 0.82 | 0.61 | 2.71 |
| Ti | 2.87 | 1.89 | 0.00 | 1.46 | 0.86 | 0.00 | 0.00 | 39.93 | 0.00 | 0.00 | 0.00 | 0.83 | 1.71 | 8.48 |
| Au | 0.00 | 0.00 | 4.29 | 0.00 | 0.00 | 3.62 | 2.59 | 0.00 | 0.00 | 0.00 | 0.00 | 0.00 | | |
| Ni | 2.46 | 0.33 | 0.84 | 0.00 | 0.00 | 0.18 | 0.14 | 0.07 | 0.00 | 0.04 | 0.09 | 0.00 | 0.03 | 0.08 |
| Cr | 0.68 | 1.60 | 0.00 | 0.00 | 0.00 | 0.00 | 0.00 | 0.00 | 0.00 | 0.01 | 0.00 | 0.02 | 0.03 | 0.02 |
| Se | 0.21 | 1.45 | 0.43 | 1.97 | 3.96 | 3.00 | 1.66 | 0.00 | 0.20 | 0.26 | 2.49 | 0.01 | | |
| Co | 0.36 | 1.37 | 0.00 | 0.01 | 0.01 | 0.06 | 0.07 | 0.05 | 0.02 | 0.06 | 0.32 | 0.02 | 0.01 | 0.01 |
| Sn | 0.04 | 0.60 | 0.00 | 0.00 | 0.00 | 0.00 | 0.00 | 0.00 | 0.00 | 0.00 | 0.00 | 0.00 | | |
| Ga | 0.24 | 0.41 | 0.00 | 0.02 | 0.00 | 0.01 | 0.02 | 0.01 | 0.00 | 0.03 | 0.00 | 0.00 | | |
| Pb | 0.64 | 0.00 | 0.97 | 0.50 | 0.51 | 0.10 | 0.11 | 0.00 | 0.13 | 0.03 | 0.00 | 0.00 | 0.20 | 0.20 |
| As | 0.04 | 0.21 | 0.02 | 0.05 | 0.00 | 0.00 | 0.06 | 0.00 | 0.00 | 0.00 | 0.00 | 0.00 | | |
| Cs | 0.18 | 0.40 | 0.06 | 0.07 | 0.08 | 0.07 | 0.06 | 0.06 | 0.16 | 0.07 | 0.02 | 0.01 | | 0.03 |
| U | 0.12 | 0.41 | 0.00 | 0.00 | 0.00 | 0.01 | 0.01 | 0.07 | 0.03 | 0.02 | 0.00 | 0.01 | 0.03 | 0.04 |
| Ce | 0.45 | 0.69 | 0.02 | 0.02 | 0.02 | 0.08 | 0.10 | 0.05 | 0.02 | 0.06 | 0.04 | 0.00 | | |
| La | 0.29 | 0.63 | 0.02 | 0.02 | 0.01 | 0.05 | 0.06 | 0.05 | 0.01 | 0.03 | 0.03 | 0.00 | | |
| Xe | 0.00 | 0.03 | 0.00 | 0.00 | 0.00 | 0.00 | 0.28 | 0.00 | 0.00 | 1.56 | 1.49 | 0.00 | | |
| Sc | 0.16 | 0.25 | 0.00 | 0.00 | 0.00 | 0.03 | 0.01 | 0.04 | 0.01 | 0.02 | 0.00 | 0.01 | | |
| Th | 0.00 | 0.26 | 0.01 | 0.00 | 0.00 | 0.01 | 0.01 | 0.17 | 0.00 | 0.00 | 0.00 | 0.00 | 0.03 | 0.04 |
| Mo | 0.00 | 0.14 | 0.00 | 0.00 | 0.00 | 0.00 | 0.00 | 0.01 | 0.00 | 0.00 | 0.00 | 0.04 | 0.06 | 0.13 |
| Sb | 0.04 | 0.28 | 0.00 | 0.00 | 0.00 | 0.00 | 0.00 | 0.01 | 0.00 | 0.01 | 0.00 | 0.00 | | |
| Tl | 0.05 | 0.10 | 0.06 | 0.00 | 0.00 | 0.08 | 0.08 | 0.00 | 0.00 | 0.00 | 0.06 | 0.03 | | |
| Be | 0.04 | 0.12 | 0.00 | 0.00 | 0.00 | 0.00 | 0.00 | 0.00 | 0.00 | 0.00 | 0.00 | 0.00 | | |

**Solid samples**

| Crystalline phases | Scores | | | |
|---|---|---|---|---|
| | **7DAL5** | **7DAL7-S** | **7YL-S3** | **7YL-w2cryst** |
| NaCl | 44 | 63 | 35 | |
| $CaPO_3$ (OH). $2H_2O$ | 8 | | | |
| NaAl ($CO_3$) $(OH)_2$ | 0 | | | |
| CaPO3 (OH) 2H2O | | 41 | | |
| $KMgCl_3$ $6H_2O$ | | | 60 | 54 |
| FeS | | | 23 | |
| $MgFe_2O_4$ | | | 20 | |
| $Na(NO_3)$ | | | 13 | |
| $TiO_2$ | | | 34 | |
| $Fe_3Si$ | | | | 25 |
| $Ca_2Mg$ $Cr_2$ $(SiO_4)$ | | | | 18 |
| FeSe | | | | 21 |

Supplementary Table 3. Chaotropicity, ionic strength and water activity for a selection of samples of the Dallol area. Chaotropicity was measured experimentally (see Methods) and also calculated, together with ionic strength values, from dominant Na, K, Mg, Ca, Fe chemistry data; water activity values were measured using a probe (see Methods). Known limits for life for each parameter are listed at the top of the table. Samples beyond that threshold for one or more of those parameters are shaded in grey.

| | | Measured chaotropicity (kJ/kg) | Calculated chaotropicity (kJ/kg) | Ionic strength (mol/L) | Water activity ($a_w$) |
|---|---|---|---|---|---|
| Life threshold* | | ≤87.3 | | ≤12.141 | ≥0.585 |
| Cave water | Gt | | n.d | n.d | 0.728 |
| | 7Gt | -18.3 | -23.80 | 4.751 | 0.729 |
| | 8Gt | -57.5 | -56.65 | 6.873 | 0.731 |
| Lake Assale | 8Ass | n.d. | 7.10 | 7.274 | 0.718 |
| Geothermally influenced Salt Plain | PS3 | n.d. | 24.09 | 7.138 | n.d |
| Dallol dome hydrothermal pools | DAL 4.00 | -21.7 | -17.87 | 6.104 | 0.719 |
| | DAL 4.0 | n.d. | -18.71 | 7.307 | n.d |
| | DAL 4A | n.d. | -9.61 | 6.346 | n.d |
| | DAL 4D | n.d. | 2.14 | 7.104 | n.d |
| | DAL 6A | n.d. | -23.97 | 7.203 | n.d |
| | DAL 9A | n.d. | -7.77 | 7.529 | n.d |
| | DAL 9C | n.d. | -16.15 | 8.349 | n.d |
| | 7DAL4-W1 | 19.3 | 40.44 | 6.314 | 0.667 |
| | 7DAL4-W2 | 8.3 | 14.28 | 5.383 | 0.698 |
| | 7DAL7 | 8.8 | 19.64 | 5.989 | 0.694 |
| | 7DAL-N1 | 9.2 | 20.84 | 6.472 | 0.694 |
| | 7DAL-N2 | 11.5 | 11.01 | 5.940 | 0.698 |
| | 7DAL9 | -8.2 | 2.95 | 5.176 | 0.708 |
| | 7DAL10 | 2.1 | -7.46 | 5.037 | 0.714 |
| | 7DAL10-1 | n.d | n.d | n.d | 0.580 |
| | 7DAL12 | -31.2 | -20.57 | 5.793 | n.d |
| | 7DAL13-W1 | -24.8 | -20.13 | 4.785 | 0.723 |
| | 7DAL14 | -11.7 | 7.54 | 5.307 | 0.748 |
| Black Lake area pools | PSBL1 | 108.3 | n.d | n.d | 0.334 |
| | PSBL2 | 93.5 | n.d | n.d | 0.345 |
| | PSBL3 | 63.4 | n.d | n.d | 0.722 |
| | PSBL4 | 61.8 | n.d | n.d | 0.711 |
| Black Lake | BL | 288.3 | 354.19 | 19.155 | 0.319 |
| | 7BL-W1 | 198.5 | 259.41 | 14.206 | 0.322 |
| | 7BL-W2 | 201.3 | 268.89 | 14.721 | n.d |
| Yellow Lake | YL1 | n.d. | 492.06 | 19.141 | n.d |
| | YL2 | n.d | 574.04 | 22.085 | n.d |
| | YL3 | 231.8 | n.d | n.d | 0.319 |
| | 7YL-W1 | 320.8 | 495.01 | 18.446 | 0.261 |
| | 7YL-W2 | 308.2 | 328.92 | 13.796 | 0.467 |
| | 7YL-W3 | n.d. | 466.64 | 17.609 | n.d |

* Data from Hallsworth et al (2007) and Stevenson et al (2015 and 2017)

Supplementary Table 4. ANOVA analysis carried out for the environmental variables and coordinates of the samples (PC1 and PC2) based on the first 2 principal components.

| | Areas (mean± SD)* | | | | | | | |
|---|---|---|---|---|---|---|---|---|
| | Dallol | Black Lake | Black Lake area | Yellow Lake | Cave water | Assale (Karum)** | F | p |
| pH | -0.45±0.22 [a] | 3.04±0.66 [b] | 3.03±0.54 [b] | 1.93±0.44 [c] | 6.36±0.16 [d] | 6.54 | 199.14 | <0.00 |
| $a_w$ | 0.703±0.23 [a] | 0.32±0.00 [b] | 0.53±0.22 [ab] | 0.35±0.11 [b] | 0.73±0.00 [a] | 0.72 | 10.97 | <0.00 |
| Temperature (°C) | 39.25±16.14 [a] | 65.30±7.49 [ab] | 40.30±0.35 [a] | 37.07±3.11 [a] | 26.47±4.39 [ac] | 21.4 | 3.59 | <0.05 |
| Salinity (% w/v) | 40.75±3.85 [a] | 76.00±0.00 [b] | 48.00±14.05 [a] | 56.67±11.55 [ab] | 30.00±0.00 [a] | 31 | 12.24 | <0.00 |
| TDS+TSS (g/l) | 393.43±35.59 [a] | 769.02±71.86 [b] | 493.72±160.09 [a] | 796.67±133.89 [b] | 326.19±4.01 [a] | 365.6 | 17.46 | <0.00 |
| PC1 | -0.424±0.26 [a] | 2.025±0.16 [b] | 0.158±0.93 [ac] | 0.772±0.6 [bc] | -1.201±0.10 [ad] | -1.077 | 18.95 | <0.00 |
| PC2 | -1.055±0.28 [a] | 0.322±0.10 [b] | 0.452±0.12 [b] | 0.366±0.11 [b] | 1.631±0.14 [c] | 1.594 | 101.51 | <0.00 |

*Different letters indicate significant differences (p<0.05) in post-hoc analysis (Bonferroni) between areas.
**Assale sample was not taken into account in the ANOVA because it only included a value.

| | | | | | | | | | | | | |
|---|---|---|---|---|---|---|---|---|---|---|---|---|
| 69 | 152298 | 146065 | 5023 | 5023 | 2 | 0.00 | 0.01 | 2 (0) | 0 | 0 | 1.00 | 0.00 |
| 53 | 126665 | 123020 | 0 | 0 | 0 | 1.00 | 0.00 | 0 (NA) | 0 | 0 | 1.00 | 0.00 |
| 34 | 124804 | 120619 | 0 | 0 | 0 | 1.00 | 0.00 | 0 (NA) | 0 | 0 | 1.00 | 0.00 |
| 68 | 233935 | 224050 | 4314 | 4303 | 21 | 0.03 | 0.03 | 35 (11) | 11 | 4 | 0.45 | 0.64 |
| 94 | 112383 | 108675 | 6 | 0 | 0 | 1.00 | 0.00 | 0 (NA) | 6 | 2 | 0.28 | 0.65 |
| 60 | 176331 | 172030 | 2 | 0 | 0 | 1.00 | 0.00 | 0 (NA) | 2 | 2 | 0.50 | 1.00 |
| 15 | 151700 | 148869 | 41 | 0 | 0 | 1.00 | 0.00 | 0 (NA) | 41 | 4 | 0.34 | 0.49 |
| 58 | 131862 | 126420 | 2 | 0 | 0 | 1.00 | 0.00 | 0 (NA) | 2 | 2 | 0.50 | 1.00 |
| 08 | 106589 | 104746 | 0 | 0 | 0 | 1.00 | 0.00 | 0 (NA) | 0 | 0 | 1.00 | 0.00 |
| 51 | 132970 | 128002 | 1 | 1 | 1 | 0.00 | NA | 1 (0) | 0 | 0 | 1.00 | 0.00 |
| 39 | 42043 | 41253 | 2 | 2 | 2 | 0.50 | 1.00 | 3 (2) | 0 | 0 | 1.00 | 0.00 |
| 16 | 158249 | 152576 | 1821 | 1 | 1 | 0.00 | NA | 1 (0) | 0 | 0 | 1.00 | 0.00 |
| 67 | 217096 | 192709 | 2 | 0 | 0 | 1.00 | 0.00 | 0 (NA) | 2 | 1 | 0.00 | NA |
| 63 | 212784 | 205528 | 1 | 1 | 1 | 0.00 | NA | 1 (0) | 0 | 0 | 1.00 | 0.00 |
| 56 | 44540 | 40224 | 62 | 0 | 0 | 1.00 | 0.00 | 0 (NA) | 62 | 2 | 0.03 | 0.12 |
| 09 | 168500 | 162187 | 2096 | 2094 | 5 | 0.01 | 0.02 | 5 (0) | 2 | 2 | 0.50 | 1.00 |
| 8 | 82170 | 71068 | 471 | 345 | 30 | 0.94 | 0.89 | 32 (3) | 126 | 7 | 0.69 | 0.72 |
| 30 | 33832 | 33711 | 45 | 31 | 5 | 0.68 | 0.82 | 5 (0) | 14 | 7 | 0.83 | 0.94 |
| 18 | 103261 | 102476 | 1492 | 1490 | 13 | 0.81 | 0.69 | 15 (3) | 2 | 2 | 0.50 | 1.00 |
| 10 | 184641 | 180701 | 5243 | 3208 | 22 | 0.91 | 0.82 | 32 (10) | 2035 | 19 | 0.84 | 0.78 |
| 99 | 130259 | 129730 | 261 | 212 | 8 | 0.72 | 0.70 | 9 (1) | 49 | 1 | 0.00 | NA |
| 25 | 100280 | 99550 | 298 | 298 | 7 | 0.66 | 0.67 | 7 (0) | 0 | 0 | 1.00 | 0.00 |
| 41 | 143552 | 142694 | 12589 | 12460 | 11 | 0.06 | 0.08 | 11 (0) | 129 | 2 | 0.48 | 0.97 |
| 74 | 226389 | 217444 | 42 | 1 | 1 | 0.00 | NA | 1 (0) | 41 | 4 | 0.30 | 0.43 |
| 84 | 177903 | 172455 | 1770 | 1302 | 13 | 0.57 | 0.47 | 13 (0) | 468 | 7 | 0.74 | 0.76 |
| 97 | 65556 | 62547 | 0 | 0 | 0 | 1.00 | 0.00 | 0 (NA) | 0 | 0 | 1.00 | 0.00 |
| 07 | 152918 | 144028 | 2 | 2 | 1 | 0.00 | NA | 1 (0) | 0 | 0 | 1.00 | 0.00 |
| 11 | 188312 | 172260 | 126 | 0 | 0 | 1.00 | 0.00 | 0 (NA) | 126 | 3 | 0.03 | 0.08 |
| 22 | 232611 | 210691 | 3 | 0 | 0 | 1.00 | 0.00 | 0 (NA) | 3 | 2 | 0.44 | 0.92 |
| 45 | 158488 | 157145 | 979 | 898 | 15 | 0.86 | 0.83 | 15 (0) | 81 | 4 | 0.07 | 0.14 |
| 58 | 86366 | 85180 | 207 | 157 | 4 | 0.60 | 0.76 | 4 (0) | 50 | 2 | 0.08 | 0.24 |
| 89 | 200588 | 191536 | 10711 | 10691 | 2 | 0.00 | 0.00 | 2 (0) | 20 | 3 | 0.27 | 0.47 |
| 32 | 123877 | 122505 | 36177 | 36016 | 39 | 0.79 | 0.50 | 39 (0) | 161 | 6 | 0.79 | 0.91 |
| 38 | 85072 | 84444 | 668 | 547 | 10 | 0.75 | 0.70 | 10 (0) | 121 | 5 | 0.71 | 0.83 |
| 95 | 76209 | 73977 | 12 | 0 | 0 | 1.00 | 0.00 | 0 (NA) | 12 | 6 | 0.78 | 0.91 |
| 66 | 227636 | 218708 | 0 | 0 | 0 | 1.00 | 0.00 | 0 (NA) | 0 | 0 | 1.00 | 0.00 |
| 97 | 177131 | 171404 | 3 | 0 | 0 | 1.00 | 0.00 | 0 (NA) | 3 | 1 | 0.00 | NA |
| 86 | 158630 | 151304 | 1 | 0 | 0 | 1.00 | 0.00 | 0 (NA) | 1 | 1 | 0.00 | NA |
| 18 | 127100 | 124137 | 8 | 0 | 0 | 1.00 | 0.00 | 0 (NA) | 8 | 4 | 0.66 | 0.88 |
| 12 | 180562 | 177286 | 5 | 3 | 2 | 0.44 | 0.92 | 2 (0) | 2 | 1 | 0.00 | NA |
| 82 | 149968 | 146028 | 146028 | 118980 | 291 | 0.88 | 0.56 | 313 (7) | 26537 | 123 | 0.96 | 0.56 |
| 90 | 303656 | 282495 | 282495 | 274013 | 602 | 0.94 | 0.51 | 701 (22) | 5893 | 39 | 0.30 | 0.51 |
| 26 | 172959 | 165299 | 165299 | 140633 | 1013 | 0.94 | 0.59 | 1190 (33) | 23963 | 496 | 0.69 | 0.59 |
| 82 | 314213 | 288299 | 288299 | 244551 | 1349 | 0.94 | 0.57 | 1441 (18) | 39873 | 859 | 0.88 | 0.57 |
| 92 | 303083 | 288086 | 288086 | 274039 | 656 | 0.87 | 0.53 | 755 (25) | 14047 | 173 | 0.91 | 0.53 |
| 33 | 71368 | 64143 | 64143 | 59788 | 524 | 0.95 | 0.57 | 735 (46) | 4353 | 147 | 0.92 | 0.57 |
| 97 | 235111 | 172010 | 172010 | 132967 | 1495 | 0.97 | 0.57 | 1565 (13) | 39039 | 227 | 0.77 | 0.57 |
| 82 | 198288 | 189428 | 32807 | 30700 | 68 | 0.71 | 0.40 | 80 (7) | 2107 | 2 | 0.00 | 0.01 |
| 63 | 93926 | 87922 | 16162 | 15575 | 64 | 0.83 | 0.50 | 71 (5) | 587 | 1 | 0.00 | NA |
| 54 | 135086 | 133629 | 202 | 136 | 16 | 0.55 | 0.50 | 19 (3) | 66 | 1 | 0.00 | NA |
| 40 | 150409 | 138429 | 14250 | 7334 | 33 | 0.64 | 0.43 | 36 (4) | 6916 | 18 | 0.64 | 0.48 |
| 71 | 105571 | 96569 | 94348 | 2876 | 19 | 0.69 | 0.49 | 26 (7) | 91472 | 351 | 0.97 | 0.69 |
| 21 | 148866 | 140175 | 135739 | 2782 | 16 | 0.83 | 0.72 | 17 (2) | 132957 | 484 | 0.97 | 0.72 |

es

| | | | | | | | | |
|---|---|---|---|---|---|---|---|---|
| 43 | 319549 | 312333 | 307451 | | 306 | 0.68 | 0.25 | 308 (2) |
| 63 | 148396 | 142049 | 140678 | | 122 | 0.65 | 0.31 | 136 (7) |
| 26 | 83207 | 82325 | 82316 | | 50 | 0.06 | 0.04 | 55 (4) |
| 71 | 87459 | 86745 | 54 | | 6 | 0.36 | 0.46 | 6 (0) |
| 63 | 14998 | 14883 | 1795 | | 4 | 0.25 | 0.34 | 4 (0) |
| 95 | 57773 | 56359 | 56269 | | 125 | 0.56 | 0.23 | 125 (0) |

**Supplementary Table 6. Mineral phases observed by SEM-EDX in precipitates of typical abiotic morphology and 'biomorphs'.** Biomorphs correspond to rounded-shaped crystalline morphs resembling cell structures (cocci, rods) and compatible with cellular sizes. Observed dominant phases are highlighted in bold

| Site | Samples | Mineral phases | |
|---|---|---|---|
| | | Typical 'crystals' | Abiotic 'Biomorphs' |
| Cave water | Gt2016, 7Gt, 8Gt_1 | Si, Ca sulfate, Fe-K sulfate, Al-Mg-Fe oxides, Fe and Ca oxides | Fe-Al silicates |
| Lake Assale (Karum) | 8Ass_2, 8Ass_3, 8Ass_4, 8Ass_6, 8Ass_7, 8Ass_8 | NaCl, Na-K-Mg chloride | **Si biomorphs** (and encrustment) |
| Dallol dome (ponds) | Dal4.0, 7DA7_07, DAL4D, 7DA9-P1, 7CA9_P1_3, 7DA7_04, 7DA7_05, 7DA7_06, 7DA9_P1_2, 7DA9_P1_5, 7DA9_P3_10, 7DA9_P3_12 | NaCl, Na-K-Mg chloride, Fe-K oxides, Ti oxides | Sulfur biomorphs, **Si biomorphs**, S-rich Na-K silicates, locally S-rich Si biomorphs, **Fe phosphates**, Fe-K phosphate, Si biomorphs – enriched in Fe, Mg, K and locally S |
| Yellow lake | YL1-03_4, 7YL_4, YL1-03_5, 7YL_6 | Fe chloride, Mg chloride | Si, $CaCl_2$, Ca phosphate |
| Black lake area (ponds) | BLPS_05_5 | Mg-Fe-K chloride | Mg chloride |
